# Tel1 is recruited at chromosomal loop/axis contact sites to regulate meiotic DNA double-strand breaks interference

**DOI:** 10.1101/2023.10.17.561996

**Authors:** Marie Dorme, Pierre Luciano, Christelle Cayrou, Rakesh Aithal, Julien Vernerey, Valérie Borde, Vincent Géli, Bertrand Llorente, Valérie Corinne Garcia

## Abstract

During meiosis, the programmed formation of DNA double-strand breaks (DSBs) by Spo11, a conserved topoisomerase II-like protein, initiates homologous recombination that leads to crossovers between homologous chromosomes, essential for accurate chromosome segregation and genome evolution. The DSB number, distribution and timing of formation are regulated during meiosis to ensure crossing over on all chromosomes and prevent genome instability. In *S. cerevisiae*, DSB interference suppresses the coincident formation of DSBs in neighboring hotspots through a Tel1/ATM dependent mechanism that remains unexplored. Here, we demonstrate that Tel1 is recruited to meiotic DSB hotspots and chromosomal axis sites in a DSB-dependent manner. This supports the tethered loop-axis complex (TLAC) model that postulates meiotic DSBs are formed within the chromosome axis environment. Tel1 recruitment to meiotic DSBs, DSB interference and the meiotic DNA damage checkpoint are all dependent on the C-terminal moiety of Xrs2, known to mediate Tel1-Xrs2 interaction in vegetative cells. However, mutation of the Xrs2 FxF/Y motif, known to stabilize Tel1 interaction with Xrs2, does not affect DSBs interference but abolishes the Tel1-dependent DNA damage checkpoint. Altogether, this work uncovers the dynamic association of Tel1 with meiotic chromosomes and highlights the critical role of its interaction with Xrs2 in fine-tuning both the meiotic DNA damage checkpoint and DSB interference.

**AUTHOR SUMMARY:** During meiosis, a special type of cell division that produces gametes, cells intentionally break their own DNA at preferred genomic sites (hotspots) to help shuffle genetic material between chromosomes. The repair process of those breaks, called homologous recombination, is crucial for genetic diversity and for ensuring chromosomes are distributed correctly to gametes. The number and location of DNA breaks are tightly controlled to prevent errors that could affect genetic stability. In the yeast model *S. cerevisiae*, a protein called Tel1 (the counterpart of ATM in mammals, a DNA damage sensor) helps limit the number of breaks per cell. Our research shows that Tel1 is recruited to DNA breaks and to specific chromosome structures (called axis), supporting the idea that DNA breaks happen within the axis environment. We also found that recruitment of Tel1 to DNA breaks depends on another protein, Xrs2. This interaction is crucial to limit further DNA breaks formation and to arrest cell cycle progression in response to DNA breaks. Overall, these findings shed light on Tel1 dynamics on meiotic chromosomes, and reveal how Tel1 and Xrs2 work together to maintain genetic stability during meiosis, a process essential for healthy reproduction and evolution.

## INTRODUCTION

During the prophase of meiosis I, the scheduled formation of DNA double-strand breaks (DSBs) by Spo11, a type II-like topoisomerase, is required for homologous recombination (HR) and crossing-overs, essential for the accurate segregation of homologous chromosomes(1–5). Meiotic DSBs are concentrated within relatively narrow and non-randomly distributed permissive regions called DSB hotspots (∼190 bp wide in *S. cerevisiae* to a few kilobases in mouse) distributed along the genome, that coincide with promoter regions in budding yeast(6–8). Meiotic DSBs arise concurrently with a structural reorganization of the chromosomes and the development of nucleoprotein axial structures. During this process, the coalescence of axis sites leads to linear arrays of chromatin loops that are anchored at their base along the axial element(9,10). In *S.cerevisiae*, this axis includes the protein Red1, Hop1, and the meiosis-specific cohesion complex comprising the Rec8 kleisin. While DSB hotspots are preferentially located in chromatin loops, HR takes place in the context of the axes, and Spo11-accessory factors essential for DSB formation (Rec114, Mer2 and Mei4) localize on the chromosomal axis. This suggests that DSB hotspots are tethered to axis sites prior of DSB formation (the “tethered loop-axis complex”, or “TLAC” model (10–12)). The formation of DSBs within the context of chromosomal axes likely provides an optimal environment for inter-homologous recombination repair(11,13). Spp1, a conserved member of the histone H3K4 methyltransferase Set1 complex, recognizes H3K4 trimethylation marks around DSB hotspots on the chromatin loops, but also interacts with the axis-associated protein Mer2, thus providing a potential mechanism for interaction of DSB hotspots with DSB proteins located on the axes(14,15). Alternatively to the TLAC model, loop extrusion mediated by cohesin, that organizes chromatin loops, has been proposed as a mechanism that dynamically brings DSB hotspots into proximity with the chromosome axis(16).

Because DSBs are precursors of meiotic recombination, their number and position have a direct impact on the distribution of genetic exchanges between parental chromosomes. Importantly, the frequency and position of meiotic DSBs is highly regulated and integrated with multiple homeostatic processes that control Spo11 activity and shape the recombination distribution over different size scales(17,18). One such pathway is the negative regulation of DSB formation by the PI3K-related checkpoint kinase Tel1/ATM. In mouse testes from ATM knock-out mice, a ∼10 fold increase in DSB numbers has been measured by quantification of Spo11-oligonucleotides(19). In drosophila and yeast, Tel1 loss also increases DSB formation and recombination(20–23). Related to the control of DSB formation, in *S. cerevisiae* Tel1 suppresses the occurrence of multiple DSBs nearby on the same chromatid (over regions up to ∼70-100kb), a process defined as DSB interference(24). As a result, the formation of two DSBs within adjacent hotspots (separated by less than ∼70-100kb) occurs less frequently than expected by chance. In the absence of Tel1, DSB interference is lost, and DSBs within adjacent hotspots can form independently, leading to the formation of measurable "double-cuts" (DCs) by physical analyses(24). Furthermore, DCs occurs far more frequently than would be expected by chance in short chromosomal domains whose borders coincide with the binding sites of chromosome axis components−thus inferred as being chromatin loops− leading to negative interference values(24). Tel1 function in preventing DSBs formation between adjacent hotspots is conserved in mammals, where the absence of ATM leads to a significant increase in deletions and duplications initiated by double-cuts formed within and between recombination hotspots(25,26).

Altogether these studies showed that Tel1/ATM negatively regulates meiotic DSB formation and controls their spatial distribution via DSB interference. DSB interference was proposed to be "reactive"(18,24), i.e. DSB dependent, in line with the known function for Tel1/ATM in response to DSB formation in vegetative cells. However, the mechanism of meiotic DSB interference remains elusive, and whether Tel1/ATM recruitment to meiotic DSBs is required to mediate DSB interference is unknown.

In humans, mutations in ATM cause ataxia telangiectasia (AT), an autosomal recessive inherited disease characterized by a strong predisposition to malignancy, cerebellar degeneration, radiosensitivity, immune deficiencies, premature ageing and sterility(27). Abundant mechanistic insight into Tel1/ATM function has been unveiled over the years, and its function in response to DSBs is well characterized. Tel1/ATM is quickly recruited to DSBs in somatic cells to trigger and coordinate −via its kinase activity on a large number of substrates(28)− various complex cellular processes such as checkpoint signaling, DNA repair, chromatin remodeling, transcription and apoptosis(28). Tel1/ATM also maintains telomere length homeostasis by promoting telomerase recruitment through phosphorylation events(29–31). Several studies have demonstrated that in somatic cells Tel1/ATM interacts with the C-terminus of Xrs2/Nbs1, a subunit of the Mre11-Rad50-Xrs2/Nbs1 complex that senses and initiates repair of DSBs and short telomeres(32–34). Yet, the activation of Tel1/ATM by Xrs2/Nbs1 is not fully understood, and the precise relationship between Xrs2-dependent recruitment of Tel1/ATM to DSBs (or telomeres) and the activation of its multiple functions (such as checkpoint control and telomeres maintenance) remains to be clarified(34).

To investigate Tel1 function during meiosis, and in particular the Tel1-dependent mechanism of DSB interference, we examined Tel1 interaction with meiotic chromosomes at specific meiotic DSB hotspots and at genome-wide level. We demonstrate that Tel1/ATM interacts genome-wide with meiotic DSB hotspots. In agreement with the preferential binding of Tel1/ATM to blocked DNA ends(35,36), this interaction is stabilized in a *SAE2* defective (*sae2Δ*) DNA end-resection deficient background, where Spo11-bound DSBs accumulate(37). Remarkably, Tel1 also localizes to both DSBs hotspots and chromosomal axis in a Spo11-catalytic dependent manner, providing evidence for interaction of Spo11-DSBs localized on the chromatin loops with the chromosomal axis, in agreement with the TLAC model(10–12) or loop extrusion mechanisms(16). Interestingly, whilst the interaction of Tel1 with meiotic DSB hotspots and axis sites is largely dependent on the C-terminus of Xrs2, DSB interference itself remains active in a *xrs2-KF* mutant that also significantly reduces Tel1 recruitment to meiotic DSBs and abolishes the DNA damage pathway triggered by unresected meiotic DSBs. Such observations indicate that a stable association of Tel1 to Spo11-DSBs is not necessary to mediate the process of DSB interference, while it is essential for the DNA damage checkpoint induced by unrepaired Spo11-DSBs. Altogether our data highlight the complex regulation of Tel1’s multiple functions and how these are fine-tuned through interaction with Xrs2, a subunit of the evolutionarily conserved Mre11-Rad50 complex.

## RESULTS

### Tel1 associates with meiotic DSB hotspots genome-wide

To explore Tel1 interaction with chromosomes during meiosis, we N-terminally FLAG-tagged Tel1 at the native locus of the *TEL1* gene (Methods). FLAG-Tel1 diploid cells progressed normally in meiosis I and II as monitored by the appearance of cells with two (MI) or four (MII) nuclei by DAPI staining (Fig. S1a), both in a wild-type and in a *SAE2* defective background (*sae2Δ*). This indicates that the FLAG tag does not impair Tel1 function in the meiotic DNA damage checkpoint activated by unresected DNA ends(38). FLAG-Tel1 protein level increased slightly (∼1.5 fold) during meiosis compared to pre-meiotic conditions (t=0) reaching a maximum at 4 h, before decreasing again (Fig. S1b,c), thus broadly following kinetics of meiotic DSB formation and repair (Fig.S1d-e). However in a *sae2Δ* where unprocessed DSBs accumulate and a Tel1-dependent checkpoint is activated(38) (Fig. S1a), Tel1 protein level increased ∼2.5-fold by 4 h, and up to 3-fold by 8 h (Fig. S1b,c). In a *sae2Δ* strain expressing catalytically dead *Spo11-Y135F*, where DSBs are not formed, the *sae2Δ*-dependent cell cycle arrest is alleviated (Fig. S1a), and Tel1 protein level remained similar to wild-type (Fig. S1b,c). Thus, in the *sae2Δ* background, unresected DSB ends activate the DNA damage checkpoint, leading to persistently elevated Tel1 protein level across meiosis.

In *S. cerevisiae*, average DSB hotspots activities vary over several orders of magnitude, the strongest hotspots being cleaved in ∼15% of the cells(8). Thus, the detection of protein interacting with meiotic DSBs is inherently limited by the frequency of hotspot cleavage. In order to determine whether Tel1 interacts with meiotic DSBs, we performed qPCR-based ChIP experiments of FLAG-Tel1 at two strong DSB hotspots (*GAT1* and *ARE1*, ranked 1^th^ and 14^st^ (8)) relative to 2 control sites (*CDC39* and *MRE11*). While no binding of FLAG-Tel1 at *GAT1* and *ARE1* was detected prior to meiotic DSB formation (time 0 h, Fig. 1a), Flag-Tel1 was transiently recruited at DSB hotspots at 3 and 4 h (Fig. 1a), coinciding with meiotic DSB formation (24). No significant enrichment was detected at 5 h (Fig. 1a). Thus, Tel1 transiently associates with meiotic DSB hotspots, in agreement with the short-lived nature of Spo11-DSBs in a wild-type background.

**Figure 1.**
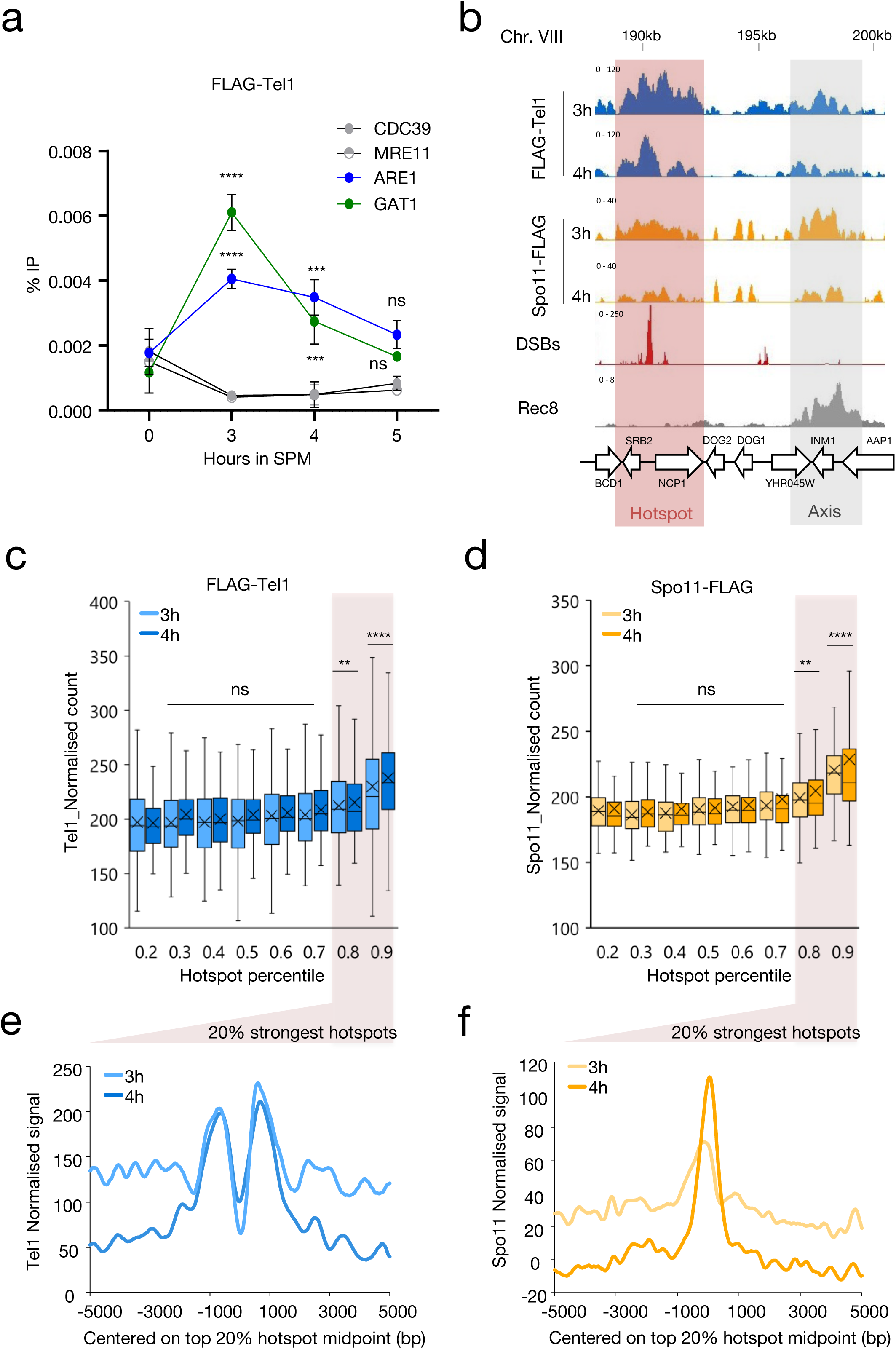
Tel1 associates with meiotic hotspots in a wild-type background. **a**, Tel1 is transiently recruited to meiotic hotspots. Mean of %IP values of FLAG-Tel1 binding, quantified by ChIP-qPCR at the *ARE1 and GAT1* hotspots relative to the *CDC39 and MRE11* control sites. Points and error bars represent mean ± SD of three independent meiotic cultures (except for time 0h for which n=2). P values are from paired ANOVA test between time 0h and all the other time, corrected via Two-stage linear step-up procedure of Benjamini, Krieger and Yekutieli for multiple comparisons test. ns (not significant); ^∗∗∗^, p < 0.001, ^∗∗∗∗^, p < 0.0001. **b**, ChIP-seq read coverage of FLAG-Tel1 (blue), Spo11-FLAG (orange) at 3 and 4 h into meiosis. A representative region of ChrVIII is shown. DSBs (red shaded area) and axis (grey shaded area) are represented by Spo11-oligos (red) and Rec8-HA ChIP-seq (grey), respectively. Horizontal axis: position on chromosome VIII. Vertical axis: Read coverage for each ChIP-seq subtracted from the input signal for Tel1 and Spo11. ChIP-seq for Rec8 (GSE69231) and Spo11 (GSE52863) as well as Spo11-oligos (GSE26449) were previously performed by Sun *et al.* and Pan J, *et al.*, respectively, in a wild-type background(8,53) **c-d**, Boxplots of Tel1 **c**) and Spo11 **d**) normalized counts per 1500bp (see Methods) partitioned by DSB strength percentile (Spo11-oligos NormHPM). P values are from Kruskal-Wallis test between percentile 0.2 and all the other percentile, corrected via Dunn’s for multiple comparisons test. ns (not significant), ^∗∗^, p < 0.01; ^∗∗∗∗^, p < 0.0001. **e**, FLAG-Tel1 read densities (RPKM per 20-bp bins) of the 8^th^ and 9^th^ percentiles ±5 kb from DSBs midpoint at 3h (light blue line) and 4h (dark blue line) in SPM, identified by Spo11-oligos(8). **f**, Spo11-FLAG read densities (RPKM per 20-bp bins) of the 8^th^ and 9^th^ percentiles ±5 kb from DSBs midpoint at 3h (light orange line) and 4h (brown line) in SPM, identified by Spo11-oligos(8).

To determine whether this localized enrichment reflects genome-wide recruitment, we performed ChIP-seq of FLAG-Tel1 at 3 h and 4 h during meiosis. At both time-points, Tel1 showed discrete ChIP-seq peaks that visually colocalize with Spo11 (39) binding sites and DSB peaks generated by sequencing of Spo11-oligos(8) (Fig. 1b, pink shaded area, Fig. S2a). A second category of Tel1 peaks also colocalized with the binding sites of Spo11-FLAG, but instead coincided with the binding sites of the axis protein Rec8(40) (Fig. 1b, grey shaded area), which define the chromosomes axis regions. Despite the visual colocalization of Tel1 and Spo11 signal with DSB hotspots at both 3 and 4h into meiosis, genome-wide analysis revealed no significant correlation with DSB hotspots strength as defined by Spo11-oligos (Fig. S2b,c). The low signal-to-noise ratio precluded reliable genome-wide peak identification and high-resolution analyses. Nevertheless, a moderate increase in Tel1 signal with hotspot strength, but still no significant correlation, emerged when considering the 20% strongest hotspots (8^th^ and 9^th^ percentiles, Fig. 1c, Fig. S2d,e r=0.42 at 3h). Similarly, an increase in Spo11 signal(39) was only observed within this subset (Fig. 1d, Fig. S2d,e r=0,48 at 3h). Even within this subset, Tel1 and Spo11 signals correlated poorly (Fig. S2e-g). Pile-up analyses further revealed that Tel1 signal at the 20% strongest hotspots displayed a bimodal distribution centered on hotspot midpoint at 3 and 4h, consistent with Tel1 binding to DSB ends that undergo resection in wild-type cells (Fig. 1e). In contrast, Spo11 signal formed a single narrower peak centered on hotspots midpoints (Fig. 1f), reflecting its catalytic role in DSB formation.

To conclude, our ChIP-qPCR experiments demonstrate that Tel1 interaction with DSBs hotspots is weak and transient during meiosis. While Tel1 and Spo11 ChIP-seq signals visually colocalize with DSB hotspots, there is no quantitative genome-wide association, likely highlighting distinct binding dynamics caused by specific functions at meiotic DSB hotspots.

### Tel1 association with meiotic DSB hotspots is stabilized in the absence of resection

Given the weak and transient association of Tel1 with meiotic DSB hotspots in wild-type cells, and prior evidence suggesting Tel1/ATM preferentially binds unresected, protein-bound DSBs (35,36), we hypothesized that Tel1 binding would be enhanced in a *sae2Δ* mutant where impaired resection preserves Spo11-DSB intermediates. In this background, Spo11 remains covalently bound to DSB ends due to the abrogation of the Mre11-dependent endonucleolytic pathway that releases Spo11 from 5’ DNA ends(37). The Tel1-dependent DNA damage checkpoint is activated(38), and only ∼50% of the cells progress through the first meiotic division at 8 h (Fig. S1a).

To investigate further Tel1 localization along chromosomes during meiosis, we assessed Tel1 binding genome-wide by ChIP-seq in the *sae2Δ* background at 4 and 6 h (experiment 1), and at 6h on 2 additional independent meiotic time-courses (experiment 2), compared to 0 h. As a control we also mapped Spo11-FLAG binding sites at 6h, in the same experimental conditions. As for FLAG-Tel1 ChIP-seq experiments, Spo11-FLAG cultures were treated with formaldehyde, thereby detecting not only covalent interactions of Spo11 with unresected DSB sites but also non-covalent interactions(41). Before the onset of meiosis, neither FLAG-Tel1 nor Spo11-FLAG were detected at specific chromosomal regions beside peaks corresponding to "hyper-ChIPable" regions that were common to both strains, and to the untagged control(42) (Fig. S3a). 4 h after meiotic entry, Tel1 shows discrete ChIP-seq peaks that increased significantly at 6 h (Fig. 2a, Fig. S3e), when the large majority of DSBs has been formed and remained unrepaired in the *sae2Δ* background (37).

**Figure 2.**
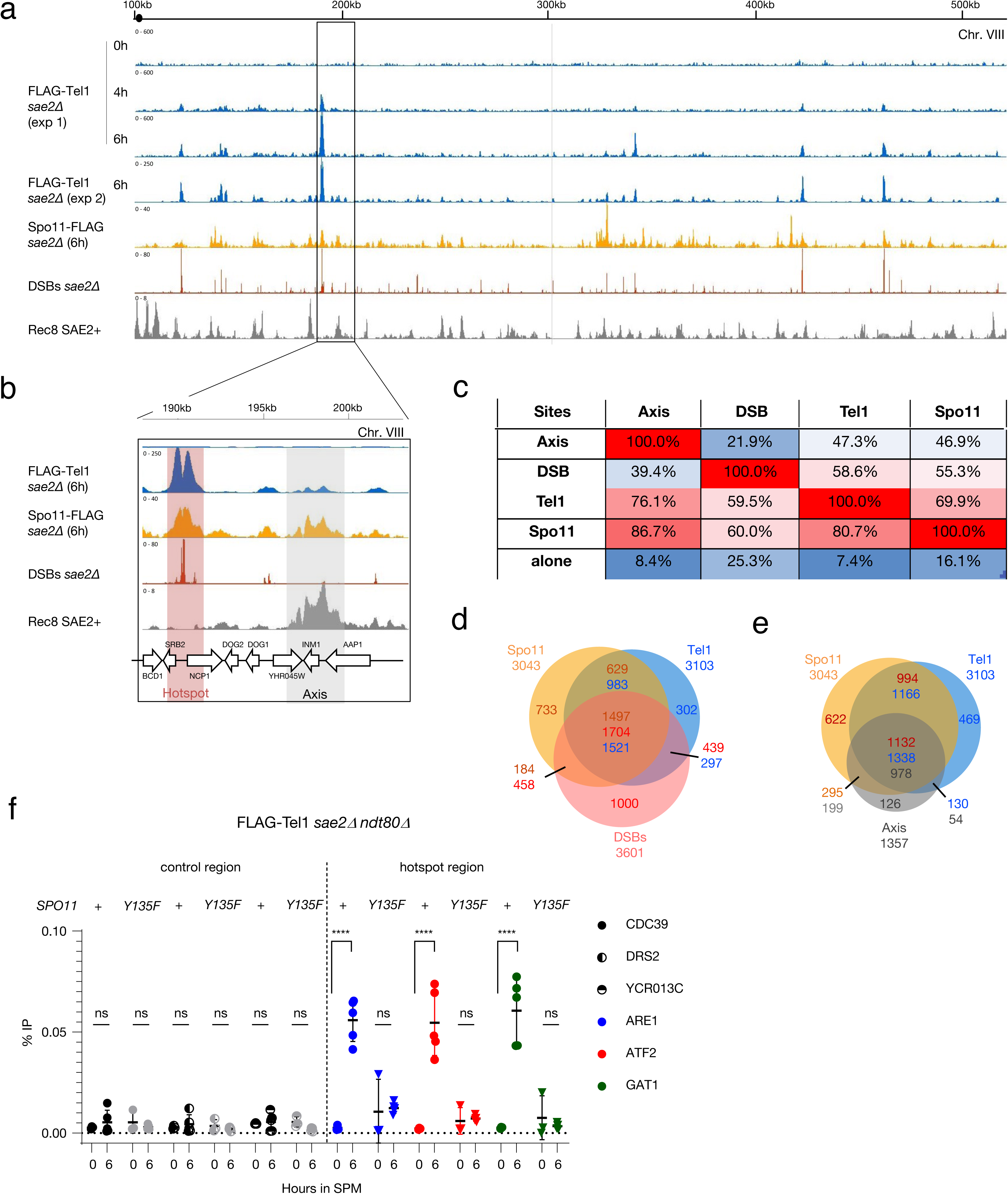
Flag-Tel1 localizes to meiotic hotspots and axis sites in *sae2Δ* background. **a**, Read coverage of FLAG-Tel1 (at 0h, 4h, 6h (exp1, n=1) and 6h (exp2, n=2)), Spo11-FLAG (n=2), DSBs (CC-seq)(43) at 6 h and Rec8-HA ChIP-seq at 4 h. Horizontal axis: position on chromosome VIII. Vertical axis: Read coverage for each ChIP-seq subtracted from the untagged signal and the signal at 0 h for Tel1 and Spo11. Tel1 and Spo11 ChIP-seq and CC-seq were performed in a *sae2Δ* background. Rec8 ChIP-seq performed in a wild-type background (GSE69231) is from Sun *et al.* (53). **b**, zoom-in on a region of ChrVIII (rectangle on panel **a**) illustrating the overlap of FLAG-Tel1 signal with Spo11-FLAG and Rec8 signals (grey shaded area), and FLAG-Tel1 signal with Spo11-FLAG and DSB sites (red shaded areas). **c**, Table of overlap between axis, DSBs, Tel1 and Spo11 peaks relative to each other’s. Percentages are indicated for the features in the first line relative to features in the first column. “Alone” in the bottom line refers to peaks overlapping with none of the other features. **d, e**, Venn diagrams showing the overlap between ChIP–seq peaks (Spo11, Tel1 and Axis) and CC-seq peaks (DSBs). Numbers are indicated using the color code of each individual feature. **f**, Tel1 recruitment to meiotic hotspots is dependent on Spo11 catalytic activity. FLAG-Tel1 recruitment measured by ChIP-qPCR at the *ARE1, ATF2 and GAT1* hotspot relative to three control sites (CDC39, DRS2 and YCR013C) in *sae2Δ ndt80Δ* and *sae2Δ ndt80Δ spo11-Y135F* mutant. Error bars indicate SD from 3 (time 0 h) and 5 (time 6 h) independent meiotic cultures. P values are from paired ANOVA test between time 0 h and 6 h, corrected via Two-stage linear step-up procedure of Benjamini, Krieger and Yekutieli for multiple comparisons test. ns (not significant), ^∗∗∗∗^, p < 0.0001.

To determine whether Tel1 and Spo11 colocalized with DSB sites, we compared ChIP-seq signals with DSBs mapped at 6 h in the *sae2Δ* background by CC-seq, a technique used to map sites of DNA–protein covalent complexes genome-wide (43). About 3,000 DSB hotspots were identified by CC-seq (43), that spatially and quantitatively correlated with DSB hotspots identified by Spo11-oligo immunoprecipitation(8,43). Consistent with the Tel1 signal at DSB hotspots in the wild-type background—but now with improved resolution—Tel1 peaks at 4 and 6h visually colocalize with both Spo11-FLAG binding sites and DSB peaks generated by CC-seq (Fig. 2b, pink shaded areas). Smoothing the 6 h FLAG-Tel1 signal with windows of increasing size revealed alternative domains of higher and lower signal that mirror peaks previously observed on DSB maps(8,44), and recapitulated by CC-seq (Fig. S3b). Similarly to ChIPs performed in wild-type cells (Fig. 1a), a second category of Tel1 peaks also colocalizes with the binding sites of Spo11-FLAG and of Rec8(40) (Fig. 2b, grey shaded area), which define the chromosomes axis regions. Tel1 and Spo11 ChIP-seq signals at 6 h correlated well between duplicates (Fig. S3c,d, r2=0.9 and 1 respectively), therefore data were pooled for subsequent finer-scale analyses. Tel1 and Spo11 peaks were determined with MACS2 and compared to each other, to DSB hotspots mapped at 6 h in the *sae2Δ* (43), and to axis coordinates (defined as the regions presents in at least two of the Red8, Hop1 and Red1 binding sites, Fig. S4a). We detected 3,103 and 3,042 peaks of Tel1 and Spo11, respectively (Fig. S4b). Aggregating signals across the genome confirmed a strong correspondence between Tel1, Spo11, DSB hotspots and axis regions (Fig. 2c-e, Fig S4c), with 81% of Tel1 peaks overlapping Spo11 peaks (p<10^-15^), 59% overlapping with DSB hotspots (p<10^-16^), and 47% with axis sites (p<10^-13^)(Fig. 2c). Tel1 peaks not colocalizing with any others features were rare (7%, Fig. 2c), and of lower intensity (p<0.0001, Fig. S5a). In agreement with previous work performed in a wild-type background(39), DSBs were depleted in axis regions (p<10^-11^, Fig. S4d) and hotspots overlapping axis sites were weaker than those overlapping Tel1 or Spo11 peaks (P<0.0001, Fig. S5b). Finally, DSB hotspots not associated with any features were of significantly low strength (P<0.0001, Fig. S5b), and visual examination showed that those hotspots often overlapped with a Tel1 signal too weak to be identified as such with our peak-calling parameters.

### Tel1 association with meiotic DSB hotspots is dependent on Spo11 catalytic activity

Mre11 is a phosphodiesterase that functions as part of the Mre11/Rad50/Xrs2 (MRX) complex, which acts as a key sensor of DSBs and facilitates DNA repair through its coordinated nuclease activities(45). While involved in the resection step of meiotic DSB ends(46), Mre11 interacts with DSB hotspots independently of DSB formation(47,48) and is required for DSB formation itself(49,50). To ask if Tel1 association with DSB hotspots strictly depends on Spo11-DSBs, or if Tel1 may have functions upstream DSB formation (like Mre11 does), we tested its chromatin binding in the catalytically dead *spo11-Y135F* mutant. Because of the precocious cell-cycle progression caused by the absence of DSBs in the *sae2Δ spo11-Y135F* mutant, and Tel1 reduced protein abundance in this background (Fig. S1b,c), ChIP experiments were performed in a *ndt80Δ* mutant background, where the cells arrest in late prophase I in a checkpoint-independent manner(51,52). In this background, where Tel1 protein level remained elevated at 6 h (Fig.S6a,b), Tel1 recruitment to *ATF2*, *ARE1* and *GAT1* hotspots was significantly reduced in the absence of Spo11 catalytic activity (Fig. 2e). This result shows that in contrast to Mre11, recruitment of Tel1 to meiotic DSB hotspots is dependent on the catalytic activity of Spo11.

### Differential association of Tel1 and Spo11 with DSB hotspots and axis regions

Our above analysis demonstrates that Tel1 peaks colocalize with Spo11 not only on DSB hotspots but also on the chromosomes axis regions (Fig. 2b-e). In order to improve our mechanistic understanding of meiotic DSB formation, we compared Tel1 and Spo11 ChIP-seq signal from the 6 h time-point (n=2) at DSB hotspots and axis sites. Tel1 normalized signal at DSB hotspots was significantly higher than at axis sites (Fig. 3a) and the strength of Tel1 signal correlated strongly with CC-seq signals (R^2^=0.72, *pearson* = 0.85, Fig. 3b, Fig. S7a). Thus, in the *sae2Δ* background, DSB hotspot strength is a major factor determining Tel1 association to DSB hotspots, suggesting a uniform binding affinity to any meiotic DSB formed, something we did not measure by ChIP-seq in the wild-type background. Heatmaps centered on DSB hotspots mid-points further confirmed the positive correlation between Tel1 peaks and DSB hotspots strength (Fig. S8a). Interestingly, Tel1 signal strength was not well correlated with the strength of axis sites as defined by Hop1 ChIP-seq read counts(53) (R^2^= 0.06, *pearson*=0.24, Fig. 3c, S7b, S8b). On the contrary, we observed a broader and stronger Spo11 signal at axis sites compared to DSB hotspots (Fig. 3d), in agreement with the self-assembly of the DSB machinery (including Spo11 dimers) on the chromosome axis before and independently of DSB formation(54–56). Accordingly, Spo11 signal strength correlated better with axis sites (R^2^=0.41, *pearson*=0.64, Fig. 3f, Fig. S7b, Fig. S8b) than with DSB hotspots (R^2^=0.27, *pearson*=0.51, Fig. 3e, Fig. S7a, Fig. S8a), consistent with DSB independent, non-covalent interactions of Spo11 with DSB hotspots being predominantly detected when performing ChIP-seq of Spo11 with formaldehyde cross-linking(41). Nevertheless, the strongest DSB hotspots (9^th^ percentile, Fig. S7a lower panel, Fig. S8a) do unveiled a significantly higher Spo11 signal (p<10^-4^). To note, Spo11 ChIP-seq signal-to-noise at DSB hotspots was largely increased in the *sae2Δ* background compared with experiments performed across time of DSB formation (3h to 5h) in a wild-type background(39) (Fig. S8a). Thus, likewise Tel1, transient interactions of Spo11 with DSB hotspots are enhanced and preserved in the *sae2Δ* background where HR is halted at the DSB stage. On the contrary, Spo11 recruitment to axis sites was not impacted by the absence of resection (Fig. S7b, Fig. S8b, compare *sae2Δ* signal-to-noise at 6h (*pearson* = 0.64) with *SAE2+* 3h (*pearson* = 0.78), once again in agreement with Spo11 assembly on the chromosome axis before and independently of DSB formation and resection.

**Figure 3.**
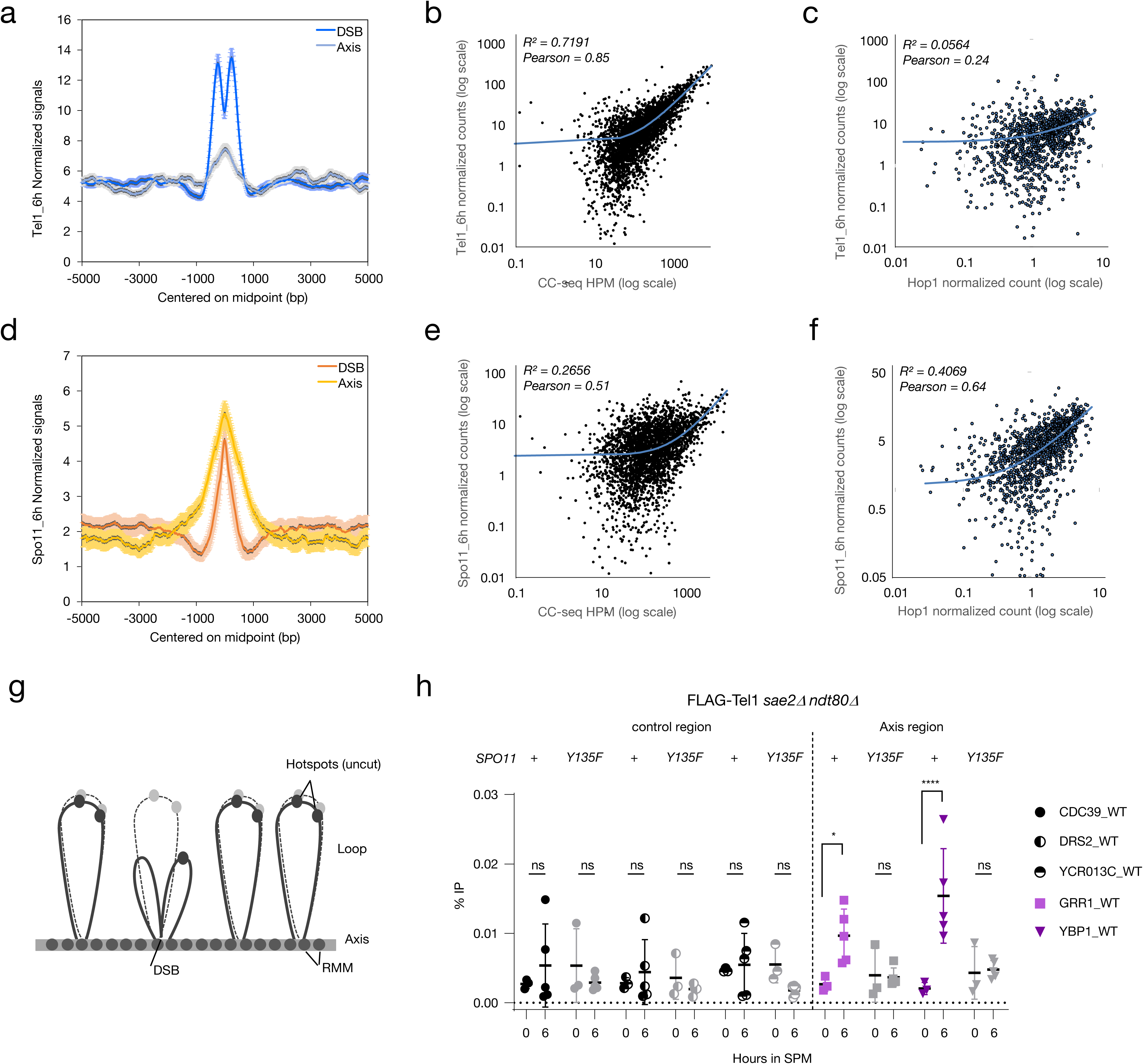
Flag-Tel1 localisation to axis sites is dependent on DSB formation. **a**, FLAG-Tel1 read densities (RPKM per 20-bp bins) ±5 kb from DSBs (blue line) and axis (grey line) midpoint, identified respectively by CC-seq(43), and by the intersect of Rec8, Red1 and Hop1 ChIP-seq (Fig. S4a)(53). **b**, Scatter plot between Tel1 signals (mean RPKM per 20-bp bins) quantified over 600pb regions centered on hotspots (93% of hotspots are <600bp), and DSB signals (CC-seq - HPM: hits per million mapped reads). **c**, Scatter plot between Tel1 signals (mean RPKM per 20-bp bins) quantified over 2kb regions centered on axis midpoint, and Hop1 signals (see methods - Read density maps). **d**, Spo11-FLAG read densities (RPKM per 20-bp bins) ±5 kb from DSBs (red line) and axis (yellow line) midpoint. **e**, Scatter plot between Spo11 signals (mean RPKM per 20-bp bins) quantified over 600pb regions centered on hotspots, and DSB signals (CC-seq - HPM). **f**, Scatter plot between Spo11 signals (mean RPKM per 20-bp bins) quantified over 2kb regions centered on axis midpoint, and Hop1 signals. The coefficient of determination (R^2^) and the Pearson correlation coefficient (Pearson) were measured and mentioned for each Scatter plot. **g**, The tethered loop-axis model for DSB formation (’TLAC’). DSB formation occurs when DSB hotspots localised in chromatin loops are tethered to axis sites(10–12). **h**, Tel1 recruitment to axis sites is dependent on Spo11 catalytic activity. FLAG-Tel1 recruitment measured by ChIP-qPCR at the *GRR1 and YBP1* axis sites relative to three control sites (*CDC39*, *DRS2* and *YCR013C)* in *sae2Δ ndt80Δ* and *sae2Δ ndt80Δ spo11-Y135F* strains. Error bars indicate SD from 3 (time 0 h) and 5 (time 6 h) independent meiotic cultures. P values are from paired ANOVA test between time 0 h and 6 h, corrected via Two-stage linear step-up procedure of Benjamini, Krieger and Yekutieli for multiple comparisons test. ns (not significant), ^∗^, p < 0.05; ^∗∗∗∗^, p < 0.0001.

### Tel1 association to axis sites is dependent on Spo11 catalytic activity

The stronger correlation of Tel1 signal with DSB hotspots than with axis sites is compatible with Tel1 being directly recruited to Spo11-DSBs, and Tel1 interaction with axis sites being indirect, caused by the spatial proximity of DSBs with axis sites mediated by loop tethering(10–12) or extrusion(16) (Fig. 3g). To verify this hypothesis, we performed ChIP-qPCR of FLAG-Tel1 at two axis sites, *GRR1* and *YBP1*, *sae2Δ ndt80Δ* and *sae2Δ ndt80Δ spo11-Y135F* mutant backgrounds. Tel1 recruitment to axis sites could be detected in the *sae2Δ ndt80Δ* background (Fig. 3h), albeit with lower efficiency compared to DSB hotspots−in agreement with differences between signals strength at DSB hotspots and axis sites already observed by ChIP-seq (Fig. 3a). Importantly, no significant recruitment of Tel1 at both axis sites was measured in the *spo11-Y135F* mutant (Fig. 3h). Thus, Tel1 association with axis sites requires Spo11-DSBs formation. Altogether, these results are consistent with a model in which Tel1 binding to the chromosomal axis is mediated by the formation of Spo11-DSBs within the context of the TLAC(10–12) or loop extrusion(16), bringing DSB hotspots, Spo11, Tel1 and the chromosome axis in close proximity.

### Xrs2 C-terminus is required for Tel1 recruitment to meiotic DSBs and activation of the DNA damage checkpoint

In mammalian cells and Xenopus extracts, Nbs1 C-terminus interacts with ATM and was proposed to be necessary for ATM recruitment to DSBs and activation(33,34,57). In *S. cerevisiae*, Tel1 recruitment to mitotic DSBs and activation of its kinase activity are abolished in a *xrs2-11* mutant, where Xrs2 is truncated for 162 amino-acids at its C-terminus(32) (Fig. 4a). We deleted the corresponding amino-acids of *XRS2* in the SK1 background to determine its impact on Tel1 recruitment to meiotic DSB hotspots and on the meiotic DNA damage checkpoint. Another Xrs2 C-terminal truncation mutant (*xrs2-664*) was previously shown to suppress the meiotic checkpoint activated by unresected DSB ends(58). In the *sae2Δ xrs2-11* mutant, similarly to the *sae2Δ tel1Δ,* approximately 80% of cells progressed through the first and second meiotic divisions by 8 h showing that the meiotic DNA damage checkpoint is abrogated (Fig. 4b). Thus, in meiotic cells, activation of the DNA-damage checkpoint by unprocessed DSBs requires the Xrs2 C-terminal domain, presumably because of its function in mediating Tel1 recruitment to DSBs.

**Figure 4.**
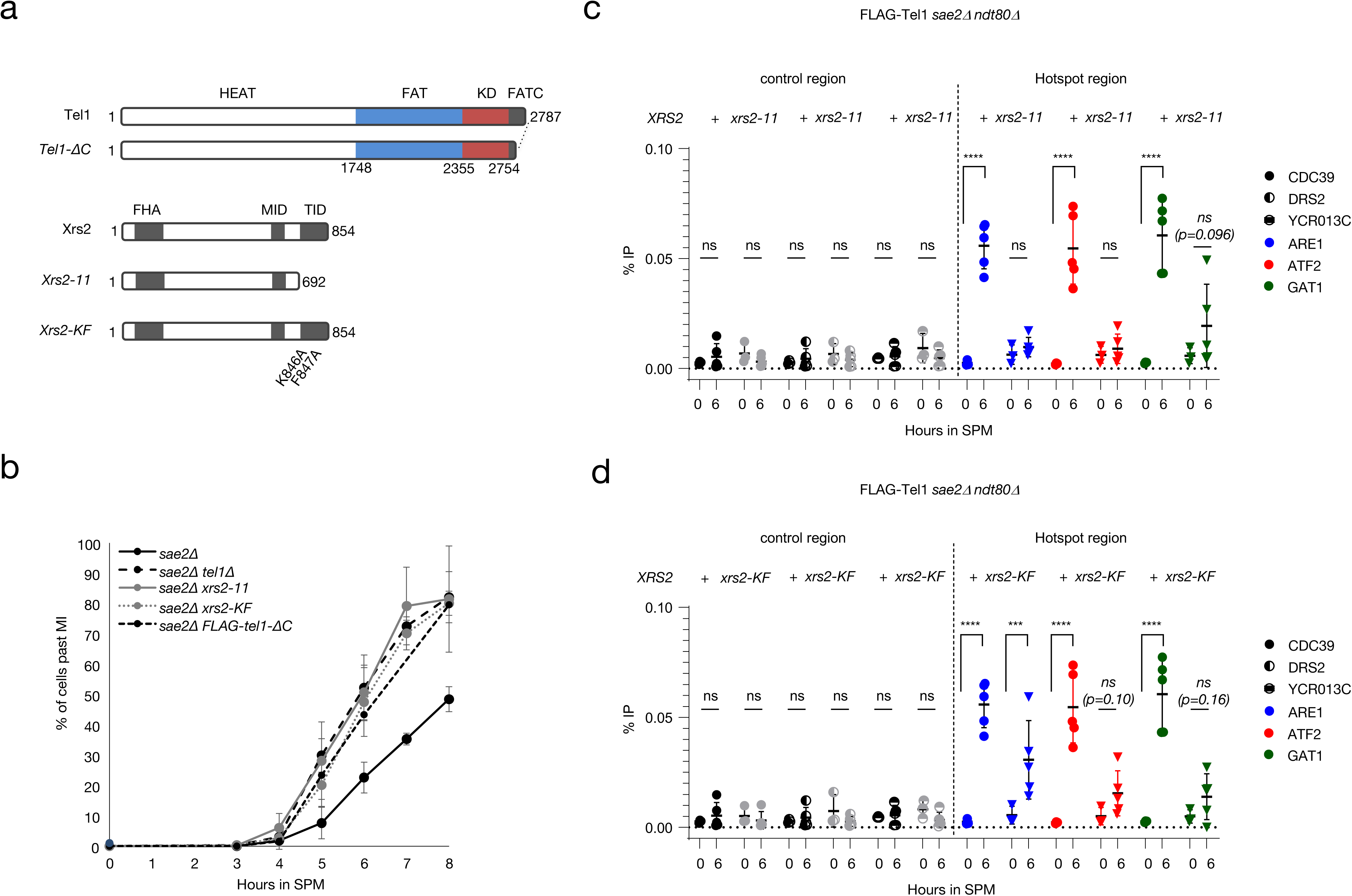
Tel1 association to meiotic DSBs is dependent on Xrs2 C-terminal domain. **a**, Schematic representation of Tel1, Xrs2 and mutants generated in this study: *xrs2-11*, *xrs2-K846A/F847A* and *tel1-ΔC*. The HEAT (Huntington’s elongation factor 3, a subunit of protein phosphatase 2A, TOR1) repeats, FAT (FRAP/ATM/TRRAP) domain, FATC (FAT C-terminal) domain of Tel1 and FHA (forkhead-associated), MID (Mre11 interaction) and TID (Tel1 interaction) domains of Xrs2, are indicated. **b**, Meiotic progression monitored by DAPI staining. The percentage of cells with 2 (MI) or 4(MII) nuclei at indicated times after entering meiosis is indicated. About 200 cells were scored for each time point. **c,d**, Tel1 recruitment to meiotic hotspots is partially dependent on Xrs2. FLAG-Tel1 recruitment measured by ChIP-qPCR at the *ARE1, ATF2 and GAT1* hotspots relative to three control sites (*CDC39*, *DRS2* and *YCR013C)* in *sae2Δ ndt80Δ* and *sae2Δ ndt80Δ xrs2-11* strains (c) and *sae2Δ ndt80Δ xrs2-K/F* strains (d). Error bars indicate SD from 3 (time 0 h) and 5 (time 6 h) independent experiments. The same %IP values are presented in c and for the *sae2Δ ndt80Δ* control. P values are from paired ANOVA test between time 0h and 6h, corrected via Two-stage linear step-up procedure of Benjamini, Krieger and Yekutieli for multiple comparisons test. ns (not significant), ^∗∗∗^, p < 0.001; ^∗∗∗∗^, p < 0.0001. P value ns but close to the significance are indicated.

To test whether Tel recruitment to meiotic DSB hotspots requires Xrs2-C terminal domain, we performed ChIP-qPCR of FLAG-Tel1 at *ARE1*, *ATF2* and *GAT1* hotspots at 6 h in the *sae2Δ xrs2-11 ndt80Δ* mutant background, to prevent precocious meiosis I exit. In this background, Tel1 protein abundance was maintained at a similar level to the *sae2Δ ndt80Δ* control (Fig. S6a,b). Importantly, no significant enrichment of Tel1 to DSBs hotspots was measured in the *sae2Δ ndt80Δ xrs2-11* mutant (Fig. 4c). We thus conclude that Xrs2 C-terminus mediates Tel1 recruitment to meiotic DSBs. Collectively, these results demonstrate that Xrs2 C-terminal domain is required for Tel1 recruitment to meiotic DSBs, which is in turn essential for activation of the meiotic DNA damage checkpoint.

The ATM-binding segment of NBS1 has been narrowed down to a sequence motif termed FxF/Y located within a conserved 20 amino-acids C-terminal domain (Fig. 4a)(33,59). In *S.cerevisiae*, mutation of the corresponding residues (K846A/F847A) reduced Xrs2 interaction with Tel1, and partially affected telomere length(59), but its impact on Tel1 recruitment to telomeres or DSBs has not been investigated(59). We introduced the corresponding mutations at the endogenous *XRS2* locus with and without the N-terminal FLAG-tag and measured meiotic progression and Tel1 recruitment to meiotic DSB hotspots. Similarly to *tel1Δ or xrs2-11* cells, the *xrs2-K846A/F847A* (*xrs2-KF*) mutant abolished the meiotic DNA-damage checkpoint induced by unresected DNA ends in the *sae2Δ* background (Fig. 4b). No significant recruitment of FLAG-Tel1 to *ATF2* and *GAT1* was measured in the *sae2Δ xrs2-KF ndt80Δ* mutant compared to the *sae2Δ ndt80Δ* control (Fig. 4d). To note however, we measured a weak but significant enrichment at the *ARE1* hotspot (Fig. 4d), indicating that Tel1 may still associates−albeit weakly−with DSB ends in the *sae2Δ xrs2-KF* mutant background.

Given the low frequency of DSB formation at meiotic DSB hotspots in the cell population, direct comparisons between mutants is challenging. Therefore, we asked if Tel1 recruitment to an HO-induced break was similarly impacted by *xrs2-11* and *xrs2-KF* mutants. To this end, we introduced FLAG-Tel1 and *XRS2* mutants in JKM139 strain, where expression of the HO endonuclease is placed under the control of a galactose-inducible promoter leading to generation of a single DSB at the MAT locus, that cannot be repaired by HR because the homologous donor loci HML or HMR are deleted(60). Tel1 binding was measured before, 90 min and 180 min after HO induction, in close proximity (0.15kb) to the DSB site and 10kb away (Fig. S9). In this system, as expected, Tel1 recruitment was more easily detected, with ∼10 fold higher %IP values (0.35-1% at 3 h) than those obtained at strong hotspots in *sae2Δ* meiotic cells (0.04-0.08% at 6h). Surprisingly, while the *xrs2-11* mutation totally abolished Tel1 recruitment as previously reported(32), the *xrs2-K846/F847* mutation had only a modest, non-significant impact on Tel1 recruitment to HO-induced breaks (Fig. S9b). Thus, unlike unresected Spo11-DSBs where Tel1 recruitment is significantly reduced in the *xrs2-K846/F847* mutant, Tel1 recruitment to HO-induced breaks is only mildly affected.

Tel1 interaction with Xrs2 also requires the C-terminal half of Tel1 FATC domain(61), which is specifically conserved among PIKKs(62). Deletion of the last nine amino-acids of the FATC domain at the extreme C-terminus of Tel1 was reported to cause defects in both DNA damage response and telomere homeostasis, and to abolish Tel1 recruitment to HO breaks(61). We deleted this short stretch of amino-acids to make the *tel1-ΔC* mutant (aa 2778 to 2787, Fig. 4a). Surprisingly, Tel1-ΔC protein level was already diminished under premeiotic conditions and failed to increase during meiosis, even in the presence of the *ndt80Δ* mutation (Fig. S6a,b). Similarly to the *sae2Δ tel1Δ*, *sae2Δ xrs2-11* and *sae2Δ xrs2-KF* mutants, the *tel1-ΔC* mutation abolished the delayed nuclear division observed in *sae2Δ* cells (Fig. 4b) and Tel1 recruitment to meiotic DSB hotspots was markedly reduced (Fig. S6c). Whether the FATC domain is directly required for Tel1 association with meiotic DSBs remains unclear, as the pronounced reduction in Tel1 binding may primarily result from the substantial decrease in Tel1 protein abundance in the *tel1-ΔC* mutant, thereby confounding results interpretation (see discussion).

### Xrs2 C-terminal domain is required for Tel1-mediated DSB interference

Meiotic DSB interference is controlled by Tel1 and prevents the formation of coincident DSBs on the same DNA molecule(24). DSB interference can be measured by physical analysis: positive interference values indicate DSB interference (i.e. coincident DSBs occur less frequently than if they were independent of each other’s), values close to zero indicate no DSB interference (i.e. independence), and negative values manifest concerted DSB formation(24). In the absence of Tel1, interference is lost and coincident cutting is observed between hotspots separated by up to 70-100kb (medium-range interference loss)(24). Coincident DSBs are also formed in close-by, adjacent hotspots far more frequently than if they were occurring independently of each other, leading to negative interference values. For example negative interference can be measured within the *HIS4::LEU2* locus (sites I and II, Fig. 5a), where concerted DSBs in the absence of Tel1 lead to abundant ∼2.4kb double-cuts hardly detected in control cells(24).

**Figure 5.**
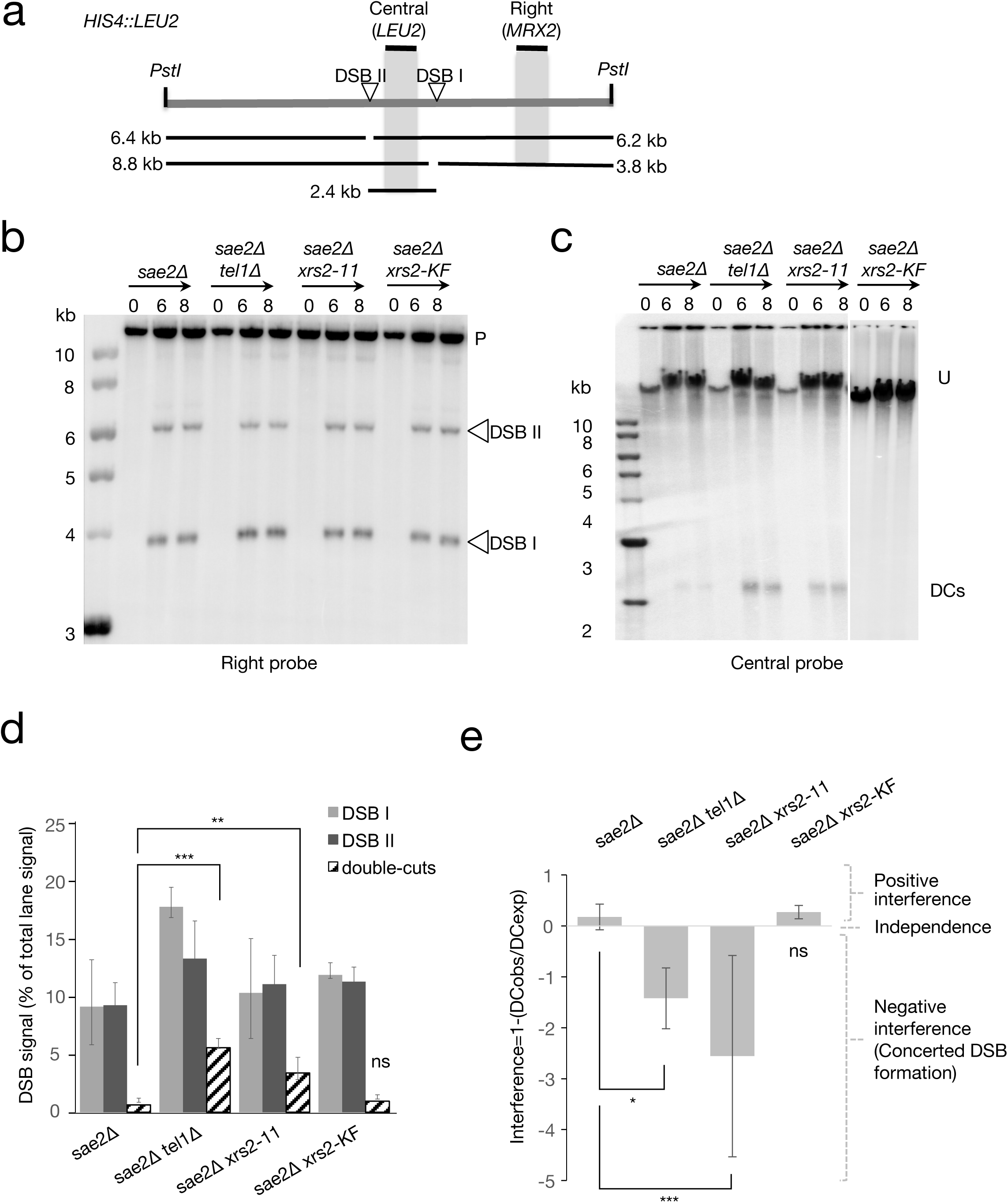
DSB interference is abolished in *xrs2-11*, but not in the *xrs2-K846/F847* (KF) mutant. **a**, Diagram of *HIS4::LEU2* locus showing DSB sites I and II positions, fragment sizes and probes for assaying DSBs and double-cuts (DCs) frequencies. **b**, **c**, PstI-digested (**b**) or undigested (**c**) genomic DNA isolated from the indicated time-points after induction of meiosis was fractionated by electrophoresis, transferred to nylon membrane and hybridized with probes as indicated (Right probe: *MRX2*, Central probe: *LEU2*). P, PstI-digested parental DNA; U, uncut parental DNA. DSB signals are marked with open triangles. Double-cuts: DCs. Parts of a unique membrane were cropped for presentation purpose. **d**, Quantification of DSBs and double-cuts (DCs) signal in b,c (average 6 and 8 h, n=3, mean +-SEM). **e**, Interference values (I=1-(observedDC/expectedDC_exp_), see methods)

To test if Tel1 recruitment to meiotic DSBs by Xrs2 is required to mediate interference, we prepared genomic DNA from *sae2Δ, sae2Δ tel1Δ* and *sae2Δ xrs2-11* mutant meiotic time-courses, and measured frequencies of DSBs at sites I and II within the *HIS4::LEU2* locus (Fig. 5b,d), as well as frequencies of double-cuts between sites I and II on undigested genomic DNA (Fig. 5c,d). Similarly to the absence of Tel1, the frequency of double-cuts was elevated in the *sae2Δ xrs2-11* mutant compared to the *sae2Δ* control (Fig. 5c,d), leading to negative values of DSB interference (Fig. 5e). Thus, the *xrs2-11* mutant phenocopies the loss of Tel1. We also probed the interval between *leu2::hisG* and *HIS4::LEU2* on chromosome 3 in order to determine if medium-range DSB interference was also lost in the *xrs2-11* mutant. Double-cuts were increased in the *xrs2-11* mutant (Fig. S10a,b), indicating a loss of DSB interference at this range also. Finally, in agreement with the severe reduction in Tel1 protein level in this mutant (Fig. S7a,b), both DSBs and double-cuts were increased in the *sae2Δ tel1-ΔC* mutant, leading to negative values of DSB interference (Fig. S11a-d). Thus, both the *xrs2-11* mutant and the *tel1-*Δ*C* mutant phenocopy a *tel1* null, conferring defects in DSB interference.

Altogether, and at first sight, these findings support the idea that Tel1 recruitment to meiotic DSB hotspots, through its interaction with Xrs2, is required to both mediate DSB interference and activate the meiotic DNA damage checkpoint. Surprisingly however, double-cuts were barely detectable in the *sae2Δ xrsK846A/F847A* mutant (Fig. 5a-d), and DSB interference was retained, with values similar to the *sae2Δ* control (Fig. 5e). Therefore, while mutations in the FxF/Y motif significantly reduce Tel1 recruitment to DSBs hotspots and abolish the Tel1-dependent meiotic DNA damage checkpoint, these residues are not required for Tel1-mediated DSB interference. This unexpected result where Tel1-mediated DSB interference requires a region in Xrs2 C-terminal domain, but not the FxF/Y motif—a motif that when mutated severely reduced but not abolished Tel1 binding to meiotic DSBs—nuances the simple “reactive” model where Tel1 recruitment to DSB ends via the Xrs2 C-terminus is required for DSB interference (see discussion). We propose that the low amount of Tel1 at meiotic DSBs measured in the *xrsK846A/F847A* mutant, insufficient to sustain the DNA damage checkpoint, may be sufficient to mediate interference.

## DISCUSSION

Tel1/ATM kinase functions and targets phosphorylation in response to DNA damages, in particular DSBs, have been extensively characterized. Although Tel1/ATM was also shown to be required in various aspects of meiotic DSBs responses, no work thus far addressed Tel1 dynamics on meiotic chromosomes, if Tel1 recruitment to meiotic DSBs is required for its known meiotic functions−such as DNA damage checkpoint activation and DSB interference.

### Tel1 protein level varies in a resection- and meiotic progression-dependent manner

We first show that steady-state Tel1 protein slightly varies during meiosis, roughly following meiotic DSB formation and repair. We can distinguish 2 phases in the variation of Tel1 protein level: an increase concomitant with the window of DSB formation (from 2 to 4 h in a wild-type background), followed by a slight decrease when DSBs are repaired (from 4 to 6 h) and a sharper drop when cells exit meiotic prophase (between 6 and 8 h). Thus, Tel1 protein abundance aligns with that of its known meiotic substrates Rec114 and Hop1(21,63), possibly reflecting a coordinated regulation during meiosis. In vegetative cells, Tel1 kinase activity is activated in response to DNA damage through post-translational modifications, recruitment to damage sites, and interactions with sensor complexes. To our knowledge, dynamic changes of Tel1 protein level in response to DNA damage has not been reported in vegetative cells. The mechanisms regulating Tel1 protein abundance in meiosis (transcriptional or post-transcriptional), and whether increased Tel1 level correlates with enhanced kinase activity, remain to be elucidated. Tel1 downregulation at later time-point, when cells exit meiotic prophase in the wild-type, may be caused by proteolysis through the 26S proteasome, a major degradation system during meiotic divisions(64).

In *sae2Δ* resection-deficient meiotic cells, Tel1 level is significantly increased compared to wild-type, and remains elevated at later time-points (8 h). Because this increase is abolished in the absence of DSB formation or in mutants that abolish the Tel1-dependent DNA damage checkpoint, we presume that both the accumulation of unresected Spo11-DNA ends and the activation of the DNA damage checkpoint contribute to steady Tel1 level in the *sae2Δ* background. Surprisingly, we observed a significant reduction in Tel1 protein level in the *tel1-ΔC* mutant compared to wild-type Tel1, whereas earlier studies reported only a slight decrease(61). The reason for this discrepancy is unclear, but may be caused by different experimental conditions or the genetic background. Tel1*-*ΔC protein level does not increase during meiosis, and given the marked reduction in protein abundance, we were unable to conclude whether this domain is required for interaction with DSB hotspots. Given that the Tel1 FATC domain was previously shown to be required for Tel1 localisation to DSB ends(61), Tel1 binding to DSBs may actively promote its stability through a positive feedback mechanism, a regulatory loop that would be disrupted in the *tel1-ΔC* mutant.

### Tel1 associates genome-wide with meiotic DSB hotspots

In order to dissect Tel1 function at meiotic DSBs and in particular in DSB interference, it was fundamental to determine in first instance whether Tel1 is recruited to meiotic DSB hotspots, and if so, whether Tel1 association to meiotic hotspots precedes, like Mre11 does(47), or requires DSB formation. In mitotic cells, Tel1 forms foci in response to γ-rays(65) and localizes to HO-induced DSBs through its interaction with the MRX complex(32). Based on these mitotic properties, Tel1 localization to meiotic DSBs was likely and anticipated. However, while HO DSBs are formed in each cell, meiotic DSBs are formed asynchronously, in at most ∼15% of the cells for the strongest hotspots at any given time, making it challenging to detect association with DSBs. Thus, although we can detect Tel1 recruitment by ChIP-qPCR in a wild-type background at strong DSB hotspots, this association is weak and transient, supporting the idea that Tel1 binds to DSB hotspots in response to DSB formation only. Consequently, correlations between Tel1 enrichment and DSB strength were obscured in our genome-wide analysis, with only the most active hotspots exhibiting detectable Tel1 recruitment.

As expected, Tel1 signal at DSB hotspots is significantly increased in the *sae2Δ* background at 4 h, and increased further at 6 h, consistent with the accumulation of unresected DSBs in this background, and with the preferential binding of Tel1 to “blocked” DNA ends(35,36). Despite Tel1 recruitment in the *sae2Δ* background being unphysiological, the higher Tel1 signal-to-noise ratio in the *sae2Δ* background enables more detailed and quantitative analyses of Tel1 localization and dynamics at DSB hotspots, and demonstrates an excellent correlation between Tel1 signal strength and DSB hotspots strength, something that could not be measured in the wild-type.

In both wild-type and *sae2Δ* background Tel1 form a bimodal signal around DSB hotspot centres, while Spo11 signal forms a single peak on DSB hotspots. This difference can be explained by the distinct roles and recruitment mechanisms of these proteins. Spo11 single peak around DSBs hotspots, reflects its direct involvement in DSB catalysis, while Tel1 recruitment to the regions flanking DSBs is consistent with its role in sensing DSBs formed by Spo11. However, and interestingly, Tel1 signal in the wild-type is broader than in the *sae2Δ* background (∼4kb versus ∼2kb, Fig. S3f), suggesting that Tel1 also binds to resected single-strand DNA ends.

Although Spo11 is likewise enriched at DSB hotspots and its ChIP-seq signal is also markedly improved in the *sae2Δ* background, its signal remains lower than that of Tel1, and the correlation between Spo11 binding and hotspot strength is correspondingly weaker. This discrepancy is also consistent with Tel1 interaction with DSB hotspots being dependent on Spo11 catalytic activity, while Spo11 was shown to interact non-covalently with DSBs hotspots independently of DSB formation (41). To note, Spo11 covalent binding to DSBs in a *sae2Δ* background may prevents adapters ligation during DNA sequencing library preparation, and favor detection of non-covalent interactions. However, this possible technical limitation should impact similarly both Tel1 and Spo11 ChiP-seq libraries preparation.

### Tel1 associates genome-wide with chromosomal axes

Meiotic DSB hotspots are located on chromatin loops, with cohesins positioned at their base, on the chromosome axes(10). Several DSB proteins, including Spo11 itself, essential for DSB formation, are also located on chromosome axes(10,12,41,54), leading to a model where DSB hotspots sequences interact with axis sites prior to DSB formation (TLAC). This model has been proposed to be critical for DSB formation, also establishing a proximity with the homologous chromosome ensuring the interhomolog bias during DSB repair(12,13). DSB-dependent interaction of Tel1 with axis sites provides direct support for the TLAC model or to alternative, non-exclusive models such as loop extrusion(16). It further suggests that broken DNA ends are not immediately released from the axis when DSBs are formed. Furthermore, our ChIP-seq data reveal localization of Tel1 signal on axis regions at both 3 and 4h (Fig.1b), indicating that DNA end resection may take place within the axis-associated chromatin environment. Tel1 localization to the axis may be crucial for phosphorylation of various targets, such as Rec114 and Hop1 that are required respectively for downregulation of DSB formation(21) and for the homologous bias(63,66). It will be particularly relevant to determine whether Tel1 interaction with the axis is required for those phosphorylation events.

While Tel1 enrichment by ChIP at DSB hotspots is significantly higher than at axis sites, Spo11 signal correlates better with axis sites. This observation is in agreement with Spo11 binding to axis sites independently of DSB formation(14,54), reinforcing the “platform” model where multiple Spo11 dimers assemble with other pro-DSB factors into DNA-driven condensates and catalyze DSB formation upon loop tethering to the axes sites(55,56).

Finally, ChIP-seq analysis in the *sae2Δ* background at the 6-hour timepoint captures proteins associated with DSBs hotspots and axis regions at a stage when breaks remain unrepaired and unresected. DSB mapping showed a strong overlap in both the localization and strength of hotspots between wild-type and *sae2Δ* cells(43,67). However, the persistence of unresected DSBs in the *sae2Δ* mutant leads to activation of the meiotic recombination checkpoint, blocking meiotic progression that may also in turn impact DSB formation, potentially affecting the interpretation of our results. Thus, while performing ChIP-seq in a *sae2Δ* is valuable for detecting otherwise transient interactions with DSBs in the wild-type, it primarily provides a static snapshot rather than reflecting the full dynamics of DSB processing and repair.

### Tel1 association with meiotic DSB hotspots and Tel1-dependent DSB interference require Xrs2 C-terminal domain

MRX binding to sites of damage precedes Tel1 localization, and Tel1–Xrs2 interaction is required for Tel1 recruitment to DSBs in vegetative cells(32,33,57,65). Tel1 binding to meiotic DSB hotspots is also dependent on Xrs2 C-terminal region, indicating that Tel1 association to Spo11-DSBs is essential for Tel1 multiple functions. In agreement with this hypothesis, both the DNA damage meiotic checkpoint and DSB interference are lost in the *xrs2-11* mutant, leading to the proposal that Tel1 recruitment to meiotic DSBs is required for both functions. Interestingly, the *xrs2-K846/F847* mutant behaves like a null mutation with regard to the meiotic DNA damage checkpoint, but Tel1 recruitment to meiotic DSBs remains significant at one of the three hotspots tested by ChIP-qPCR, and DSB interference is maintained. In agreement with its function being only partially impacted, the *xrs2-K846/F847* mutant was reported to behave almost like a null mutant regarding the DNA damage response in vegetative cells, but to cause only a modest defect in telomeres length homeostasis(59). Tel1 recruitment to telomeres in a *xrs2-K846/F847* mutant was not investigated(59), but Xrs2 interaction with Tel1 was not completely abolished (tested with an Xrs2 peptide bearing the *xrs2-K846/F847* mutation), suggesting that residual binding and recruitment of Tel1 may occur and be sufficient for some telomeric functions. In our experiments, Tel1 interaction with DSB hotspots may occur transiently in the *sae2Δ xrs2-K846/F847* mutant, more efficiently at some hotspots than others, enough to mediate DSB interference, but insufficiently to activate and maintain meiotic DNA damage signaling. In agreement with residual Tel1 interaction with Spo11-DSBs in the *xrs2-K846/F847* mutant, this mutant has only a modest, non-significant impact on Tel1 recruitment to HO-induced DSBs in vegetative condition. The differing impact of the *xrs2-K846/F847* mutation on Tel1 recruitment to HO-induced DSBs versus unresected Spo11-DSBs may stem from several factors: the inherently weaker Tel1 ChIP signal at meiotic DSBs, the contrasting dynamics between the two systems (with a single HO-induced DSB compared to up to 250 Spo11-induced DSBs per meiotic cell), and the distinct processing of these breaks—HO-induced DSBs are resected, while Spo11-DSBs remain unresected in a *sae2Δ* background. This result indicates that resection modulates Tel1 recruitment, and that the KF motif of Xrs2 motif is more critical for Tel1 association with unresected ends, a hypothesis that would requires further investigation. Altogether, we propose that Tel1 binds transiently to meiotic DSBs in the *xrs2-K846/F847* mutant, enough for catalytic activities towards specific targets and interference, but insufficiently for activation of the meiotic DNA damage checkpoint. In this scenario, Tel1-Xrs2 interaction through the FxF/Y motif would be necessary to stabilize activated Tel1 at sites of DNA damage, critical for Tel1-mediated DNA damage checkpoint activation, but not for DSB interference. In line with previous hypotheses(33), another FxF/Y motif within Xrs2 C-terminus may be required for Tel1 full recruitment and activities. To conclude, we propose that Xrs2 interaction with Tel1 at meiotic DSBs modulates Tel1 activation, with different levels of activation being required for various Tel1-dependent processes. While checkpoint activation requires robust DSBs binding (and likely strong kinase activity), DSB interference appears to be less dependent on Tel1 retention to DSBs and may also rely on kinase-independent functions.

## METHODS

### Strains and media

All *S.cerevisiae* strains used in this study are derivatives of the SK1 diploid used for its efficient meiotic entry and progression. Strain genotypes are listed in Table S1.

To generate the FLAG-Tel1 SK1 strain, a URA3 cassette was integrated upstream the *TEL1* promoter. The 5’ region of *3xFLAG-TEL1* from the FLAG-Tel1 strain generated by Seidel *et al.*(68) was amplified by PCR and transformed in the *URA3*-*TEL1* SK1 strain. After selection on 5-FOA plates, the FLAG-tag insertion was verified by PCR screening and sequencing.

Strains with deletion and mutations were constructed by one-step allele replacement. The *xrs2-11*::*kanMX6, xrs2-KF::KANMX6 and tel1ΔC::KANMX6* strains were made by transformation of a PCR product generated using pFA6a-kanMX6 and pFA6a-NatMX6 as template. Oligonucleotides are listed in table S2.

### Meiotic Time courses

Pre-culture of 8 – 10 colonies were performed in YPD + AU (Yeast Extract 1%, Peptone 2%, Glucose 2%, Adenine and Uracil 0.0137%) for 30h at 30°C, 250rpm agitation. 0.3 OD/ml of cells were incubated in 200ml pre-sporulation medium (BYTA: Yeast Extract 1%, Tryptone 2%, Potassium Acetate 1%, Potassium Hydrogen Phtalate 0.05M, Antifoam 0.001%) for 16h at 30°C with 250rpm agitation. Cells were collected and washed with water, and resuspended at 2.5 OD/ml in in pre-warmed sporulation medium (SPM: 0.3% Potassium Acetate, 0.02% Raffinose, amino acids mix and Antifoam 0.001%) before incubation at 30°C with 250rpm agitation. Sample were collected at different time point and treated accordingly to the following experiments.

### Meiotic Progression

50μl of cells were collected at different time-points and fixed in ethanol 70%. Meiotic progression was monitored by DAPI staining and counting nuclei content by fluorescence microscopy. ∼200cells were counted for each condition.

### Southern Blot

Genomic DNA was extracted from samples of synchronously sporulating cultures as previously described(69). Briefly, spheroplasts were prepared in 1M sorbitol, 0.1M EDTA, 0.1M NaHPO4 pH7.5, 1% β-mercaptoethanol and 200µg/ml zymolyase 100T for 30min at 37°C, and lysed by incubation with SDS (0.5% final) and proteinase K (200µg/ml final) for 1hour at 60°C. DNA was isolated from protein by adding equal amount of phenol:chloroform:isoamyl alcohol. DNA was precipitated by adding one-tenth volume of 3M sodium acetate pH5.2 and an equal volume of ethanol 100%. Precipitates were washed in 70% ethanol and resuspend overnight in TE 1X at 4°C. RNA was digested by RNAse A (100µg/ml final) for 60min at 37°C and DNA was reprecipitated with sodium acetate and ethanol before suspension in 1xTE buffer.

DSBs and double cuts were detected and measured as previously described(24). Briefly, to measure meiotic DSBs, DNA was digested with the appropriate restriction enzyme (see Probe Table), whereas DCs were measured on undigested DNA. Digested and undigested DNA were fractioned in 0.7% agarose – TBE 1X gel for ∼18h, transferred to nylon membrane under denaturing condition and hybridized with the appropriate probe for detection and quantification. DSBs and DCs level were quantified as previously described(24) using the Fiji Software.

### Pulse Field Gel Electrophoresis (PFGE)

DNA was prepared in plug as previously described(70). Plugs were sealed in 1% agarose – 0.5X TBE gel, and chromosomes were fractioned using a CHEF-DRIII PFGE system (Bio-Rad) using the following program: 14°C; 6 V/cm; switch angle 120°; switch time of 30 – 30s for 3 h and time 3 – 6s for 20 h. DNA was transferred to nylon membrane under denaturing condition and hybridized with a *FRM2* probe (see Table S2). DCs level were quantified by Fiji Software.

### Signal quantification and DSB interference

DSB interference values between DSB hotspots I and II at *HIS4::LEU2* were calculated as previously described(24). Briefly, double-cuts (DCobs, “observed” DC) were quantified using a central probe (*LEU2*) measuring double-cut frequency between the two hotspots. This signal was multiplied by three to take in account that this probe hybridises to three parental genomic locations (*HIS4::LEU2*, *leu2::hisG*, *nuc1::LEU2*). Expected double-cuts (DCexp) were calculated by multiplying the frequencies of DSBs measured at DSB I and II using a MRX2 probe. However, because of double-cut formation, the DSB II signal measured using the *MRX2* probe is underestimated. To compensate for this loss, the DCobs signal was added to DSB II. DSB interference were calculated as followed: *1 – DCexp / DCobs*.

### Western Blot

Proteins were isolated by trichloroacetic acid (TCA) precipitation method. 10ml of sporulating cells were treated with 250µl of TCA10% and mechanically lysed by Precellys agitation (4 cycles of 30sec at 6000rpm). TCA pellets were extracted from lysate, and dissolved in STE buffer (2% SDS, 0.5M Tris HCl pH8.1, 10mMEDTA, Bromophenol Blue) with heat at 90°C for 5min. After 3 min of centrifugation at 3000rpm at RT, whole cell supernatant was transferred in new tubes and stored at -20°C.

Protein samples were loaded on a 6% SDS-polyacrylamide gel and blotted to 0.45µm nitrocellulose membrane. Membranes were saturated with 5% BSA – PBST. FLAG-Tel1 was detected with an anti-FLAG mouse antibody (M2 monoclonal, Sigma, 1/3000) and revealed with anti-mouse (Dako P0447, 1/5000) HRP conjugated secondary antibodies. Signal was detected using the HCL kit (ECL Primer Western Blotting Detection Reagent, Amersham) and a Chemidoc system (Biorad). Signal were quantified with the ImageLab Software. Flag-Tel1 signal was normalized to ponceau staining before standardisation to time 0 h. For *ndt80* mutant experiments, Flag-Tel1 protein extracts were treated as above except that Flag-Tel1 was detected using a goat anti-mouse IgG Dylight 800 antibody (Invitrogen, SA5-10036). Flag-Tel1 signal was normalized to Rap1 protein level using a Rabbit polyclonal Rap1-antibody (Homemade, V.GELI).

### Chromatin immunoprecipitation (ChIP)

For each meiotic time point, 2 x 10^8^ cells were fixed in paraformaldehyde (1% final). FLAG-Tel1 cells were fixed for 30min, while FLAG-Spo11 strain were fixed for 15min, quenched by glycine for 5min (125mM final) and washed in TBS 1X three times. Cells were lysed using mechanical lysis (Precellys, 6 cycles at 6.5M/sec 30sec) together with a Lysis buffer (50mM HEPES-KOH, 140mM NaCl, 1mM EDTA pH8, 1% Triton X100, 0.1% Sodium Deoxycholate) complemented with protease inhibitor cocktail tablet (Roche, Ref 05 892 791 001), PMSF (1mM final), pepstatin (0.1mM final) and leupeptin (1mM final). After cell lysis, chromatin was sheared by sonication into 250 – 500pb fragment (30sec ON/30sec OFF, 7 cycles, 4°C, Diagenode Bioruptor Pico). 50µl was kept aside as the input. For Flag-Tel1 ChIP, samples were incubated with 2µl of anti-FLAG antibody (M2 monoclonal, Sigma, Ref F1804) at 4°C for 2 h 30min on a rotating wheel then 20µl of pre-equilibrated Protein-G Dynabeads (Invitrogen, 1003D) were adding and incubated for an additional 2h30min at 4°C on a rotating wheel. IP samples were washed twice with lysis buffer, twice with 500mM NaCl containing lysis buffer, twice with LiCl buffer (10mM Tris-HCl pH8, 0.25LiCl, 1mM EDTA pH8, 0.5% Triton X-100, 0.5% Sodium Deoxycholate) and once with 1XTE buffer. Chromatin was eluted from magnetic beads in elution buffer (TE 1X, 0.5% SDS) at 65°C for 10min. DNA-protein interaction was disrupted by incubation at 65°C overnight for IP and input samples. RNA was digested by RNAse A treatment (20µg total, 30min at 37°C) and proteins by proteinase K treatment (100µg total, 1hour at 42°C). DNA was purified on column (MSB Spin PCRapace, Invitek Molecular, ref: 1020220300).

To detect enrichment of proteins in proximity of HO endonuclease cut sites, cells were growth overnight in YPD at 30°C under agitation. Cells were washed once with YP-raff and incubated for an additional 5 hours in YP-raff, diluted to OD_600_ = 0.1 in YP-raff and incubated for 3 generations at 30°C. At OD_600_ = 0.8 galactose was added to the final concentration of 2% to induce expression of the HO endonuclease. At time 0, 90’ and 180’, 5.10^7^ cells were cross-linked with 1% formaldehyde for 15 min at RT, quenched for 5 min with 0.125 M Glycine, washed 3X with cold 1X TBS and pellets were snap-frozen in liquid nitrogen. ChIPs were performed as described above.

### Quantitative PCR (qPCR) on ChIP samples

qPCR was performed on ChIP samples at control sites (within *CDC39, MRE11, DRS2, YCR013C*), DSB hotspots sites (*ARE1*, *ATF2*, *GAT1*) and axis sites (*YBP1*, *GRR1*) (see qPCR Primer Table) using the TB Green Premix Ex Taq II (Tli RNAse H Plus, Takara). Each sample and input were quantified relatively to a standard curve. We calculated the percent IP value (%IP) by dividing the IP signal by the input signal. We take in account the dilution of the input for qPCR analysis (20 fold), and the difference of volume between the amount of input versus the one used for IP (10 fold) to calculate the real %IP.

### ChIP-seq

Tel1 and Spo11 ChIP-seq were performed following the ChIP-qPCR protocol with some modifications. For both FLAG-Tel1 and FLAG-Spo11 strain, cells were fixed in paraformaldehyde for 30min. Protein G magnetic beads were washed (0.1M NaHPO4 pH7.5, 0.01%Tween20, pH8.2) and saturated with BSA (0.1M NaP, 0.01% Tween20, 5mg/ml BSA) for 3 h at 4°C. Anti-FLAG antibody (M2 Monoclonal, Sigma) was pre-incubated with saturated-magnetic beads for 4 h at 4°C. Cell lysis was performed as for the ChIP-qPCR protocol. Chromatin shearing was performed in Diagenode tube 15ml with beads (900 – 1000mg) by 7 cycles of sonication (30sec ON/30sec OFF, 4°C). Chromatin was immunoprecipitated with antibody-protein G mix by overnight incubation at 4°C. Samples were washed as for the ChIP protocol, and DNA was eluted with IPure v2 Diagenode kit. DNA quantity was measured by QuBit and fragmentation quality was controlled by Agilent chip (Bio-analyser). DNA was sequenced with a NovaSeq sequencer (Illumina) using a 100nt paired-end protocol (TruSeq) at the Institut Curie sequencing facility (Paris, France).

### Data processing and computational analysis Alignment and mapping

Sequencing quality control was determined using FastQC tool (http://www.bioinformatics.babraham.ac.uk/projects/fastqc/) and aggregated across samples using MultiQC. Reads with Phred quality score less than 30 were filtered out. For Tel1 and Spo11 ChIP-seq in the *sae2Δ* mutant, paired-end reads were mapped to Cer3H4L2 *saccharomyces cerevisiae* genome (PMID: 31649282) using *Bowtie2* (v2.3.4.3)(71) with default parameters. To allow complete comparison between WT and *sae2Δ* background (concern the chrIII only), paired-end reads from Tel1 ChIP-seq in WT cells were also mapped to Cer3H4L2 *saccharomyces cerevisiae* genome (PMID: 31649282) using *Bowtie2* (v2.3.4.3)(71) with default parameters.

Only mapped reads were filtered in using *samtools view* (*--exclude-flags 4*)^67^. Duplicate tags were removed using Picard Tools *MarkDuplicates* and mapped tags were processed for further analysis. For all subsequent analyses, mitochondrial genome, plasmid genome, and ribosomal DNA (rDNA) loci were blacklisted and removed. For visualization purpose, mapped tags were converted into BedGraph and normalized by their library size (FPKM) using *bamCoverage* (72).

Wig files from Hop1, Rec8 and Red1 ChIP-seq data in the WT background were obtained from published study (PMID: 26258962; GSE69231) transformed in bedGraph file with *wig2bed* (BEDOPS v2.4.41) and converted into the Cer3H4L2 by adding 3078 bp from the position 65684 and 1173 bp bp from the position 95042 on the chrIII.

Spo11 ChIP-seq reads from GSE52863 (PMID: 24635992 - WT background; 3h, 4h and 5h after meiotic induction) and the matched untagged sequencing reads were aligned to the yeast Cer3H4L2 genome (PMID: 31649282) using *bowtie2* and duplicate sequencing reads were removed by *samtools markdup*. BedGraph files for each time point were created using *bamCoverage (*FPKM) (72).

Mre11 normalized log2 ratio from the ChIP-on-Chip published study (PMID: 21822291 - WT background; 3 h after meiotic induction; GSE30072) was transformed in bedGraph file and converted into the Cer3H4L2 by adding 3078 bp from the position 65684 and 1173 bp bp from the position 95042 on the chrIII.

For subsequent analysis, visualization, and read density maps (Fig. 1; Fig. 2a-e; Fig. 3a-f; Fig. S2; Fig. S3e-f; Fig. S5; Fig. S7 and; Fig. S8), signal (normalized counts) at each sample was first subtracted from the corresponding untagged (for the *sae2Δ*) or input signal (for the wild-type), and replicates were merged together when existing. For Tel1 and Spo11 ChIP-seq in the *sae2Δ* mutant, the signal at t 0h was further subtracted from the corresponding sample signal at t 6h before merging the replicates together.

We used Integrative Genome Viewer (IGV) v2.9.4(73,74) or Integrative Genomics Viewer (IGB – v 10.0.1)(75) for visualization.

### Correlation between biological replicates

The degree of correlation between biological replicates (Fig. S3c,d) was visualized with a scatter plot and subsequent computation of Pearson’s correlation coefficient (*FP*) and Spearman’s rank correlation coefficient (ρ) using the average signals in individual 100-bp non-overlapping genomic windows.

### Chromosome profiles

For Fig. S3b, Cer3H4L2 mapped tags were counted within 1000 bp bins (*beedtools makewindows -w 1000*)(76) throughout the genome using *featureCounts* (v1.6.4)(77). Signal values were then normalized by library size, smoothened by applying Hann function with different window sizes (2.5, 5, 10,25 and 50kb), and displayed using homemade R scripts.

### Peak calling

To ensure greater precision in peak selection, Tel1 ChIP-seq binding sites in the *sae2Δ* background at 6h were determined with *Macs2 callpeak*(78) using 3 different p-value cut-off *(--bw 300 --nomodel --keep-dup 1 -p [0.05, 1e-2 and 1e-4])* and untagged reads as the control. NarrowPeak from duplicate samples were merged for the three p-value conditions and only the smaller region from overlapping peaks was considering. For correlation analysis, regions with distance <350bp were merged and peaks of the FLAG-Tel1 *sae2Δ* t-0h peaks (“hyper-ChIPable” regions) were substracted from the FLAG-Tel1 *sae2Δ* t-6h peaks list. Spo11 ChIP-seq binding sites, in the *sae2Δ* background, were determined with *Macs2 bdgbroadcall*. As above, peaks of the FLAG-Spo11 *sae2Δ* t-0h peaks were substracted from the FLAG-Spo11 *sae2Δ* t-6h peaks list. Red1, Hop1 and Rec8 ChIP-seq binding sites in the WT background were determined with *Macs2 bdgbroadcall* (-c 1.301 -l 100 -g 100). Axis sites were obtained by cross-referencing Red1, Hop1 and Rec8 binding sites at 1 bp minimum, considering sites that overlap at least two out of the 3 proteins binding sites (Fig. S4a).

### High-ChIPable regions

The three-negative control from FLAG-Tel1 *sae2Δ,* FLAG-Spo11 *sae2Δ* and *sae2Δ* strains (see Fig. S3a) were analyzed with the Macs2 software(78) to call ChIP-seq broad peaks, subsequently merged to create the blacklist of high-ChIPable regions. 550 independent regions were found all over the genome.

For all peak calls described above, positions that overlap a hyper-ChIPable regions have been removed from the overlapping peak coordinate. Peaks > 200bp were retained.

### Intersection between dataset

Intersection between each dataset were counted as the overlaps between axis, DSBs, Tel1 and Spo11 peaks at 1 bp minimum using *intersectBed*(77). To test the significance of the intersection between datasets (real), we generated 10 sets of random peaks (shuffled). Each random set contained the same number of regions as in the cognate set of DSBs or Tel1 peaks and had the same length distribution, but each region segment was randomly picked with the condition that the segments did not overlap with any actual DSBs or Tel1 peaks (*shuffleBed* [- chrom -noOverlapping])(77).

Venn diagrams were performed with *Venn Diagram Generator* (https://academo.org/demos/venn-diagram-generator/). Boxplot were generated *by GraphPad Prism v8.0.2* (GraphPad Software, Boston, Massachusetts USA, www.graphpad.com)

### Read density maps

Read coverage plots and heatmap were obtained by extracting 5 kb on each side of the center of DSBs or axis sites and counting the reads per 20-bp bins using *annotatePeaks.pl* (HOMER)(79). The *-hist* option was used to generate histograms of position dependent read counts relative to the center of peaks (Fig. 1e,f; Fig. 3a,d) and *– ghist* to generate an output profile for each peaks used to generate heatmaps (Fig. S8). The regions were sorted according to the DSBs strength (NormHPM) for Fig. S8a and Hop1 counts (RPKM) for the Fig. S8b, and heatmaps were created using *TreeView*(80). For scatter plot of Fig. 3 and boxplot Fig. S5, the mean count per 20-bp were measured on a region of 600 bp centered on hotspot region (Fig. 3b,e, Fig. S5a) and 2000 bp centered on axis midpoint (Fig. 3c,f, Fig. S5b). For Fig. S5, DSB or Axis peaks were partitioned by strength percentile according to DSB hotspots strength (NormHPM) and Hop1 normalized count (RPKM) respectively. For scatter plot of Fig. S2b,c,f,g and boxplot Fig. 1c,d and Fig. S2d in WT strain, the read count per 1500-bp were measured on a region centered on hotspot region or the top20% Hotspot. To avoid any Tel1 or Spo11 signal due to the axis region, only regions of 1500 bp that overlap the axis by less than 10% are taken into account.

Boxplot were generated by GraphPad Prism v8.0.2 (GraphPad Software, Boston, Massachusetts USA, www.graphpad.com).

### Statistics

For qPCR, statistical analysis was performed on paired time point (0 h versus other time point) using ANOVA corrected via Two-stage linear step-up procedure of Benjamini, Krieger and Yekutieli for multiple comparisons test (alpha = 0.05).

For boxplot (Fig. 1c,d; Fig. S2d), statistical analysis was performed on unpaired data using Kruskal-Wallis test between percentile 0.2 and all the other percentile, corrected via Dunn’s for multiple comparisons test.

The significance of the difference between random (shuffled) and actual region (real) was performed using Mann Whitney test.

Error bars represent the mean ± SD of the indicated number of independent experiments. ∗p ≤ 0.05, ∗∗p ≤ 0.01, ∗∗∗p ≤ 0.001 and ∗∗∗∗p ≤ 0.0001

## DATA AVAILABILITY

Libraries are available via the GEO repository GSE276891

## FUNDING

This work was funded by the Agence Nationale de la Recherche (ANR-16-CE12-0028-01, *Meioint* project). MD phD studentship was funded by Aix-Marseille University.

## ACKNOWLEDGEMENTS

We are grateful to Matthew Neale and Christophe de la Roche Saint André for helpful discussions and constructive reading of the manuscript, Elizabeth Blackburn and Dana Smith for sending over useful strains, Samuel Granjeaud for support with statistical analyses, and Stéphane Coulon for support.

## AUTHOR CONTRIBUTIONS

VCG and BL supervised the project. MD, PL, CC, BL, VCG conceived the project. MD, PL and RA performed experiments. CC and JV performed bioinformatic analyses and statistics. VG and VB supported the project and with MD, PL, CC, BL and VCG analysed data. CC and VCG curated the data. MD and VCG wrote the manuscript.

## SUPPORTING INFORMATION

Figure S1.

**a**, Meiotic progression as the percentage of cells with 2 (MI) or 4(MII) nuclei at indicated times after entering meiosis. About 200 cells were scored by DAPI staining for each time point.

**b**, Immunoblot of meiotic TCA extracts at indicated time-points probed with anti-FLAG antibody. Bottom panels: ponceau staining.

**c**, Quantification of immunoblots (n=3) as per (b). Signal normalization was performed against ponceau staining before standardisation to time 0 h.

**d**, Southern blot of PstI digested gDNA hybridized with a probe located on the *MRX2* open-reading frame as indicated. P, parental DNA. DSB signals are marked with an asterisk.

**e**, Quantification of DSBs in d (n=1).

Figure S2.

**a**, Read coverage of FLAG-Tel1 (blue), Spo11-FLAG (orange) at 3 and 4 h in a large region of ChrVIII. DSBs are represented by Spo11-oligos (red) and axis by Rec8-HA ChIP-seq (grey). Horizontal axis: position on chromosome VIII. Vertical axis: Read coverage for each ChIP-seq subtracted from the input signal for Tel1 and Spo11. ChIP-seq for Tel1 was performed in a wild-type background. ChIP-seq for Rec8 (GSE69231) and Spo11 (GSE52863) as well as Spo11-oligos (GSE26449) were previously performed by Sun *et al.* and Pan J, *et al.*, respectively, also in a wild-type background(8,53).

**b**, Scatter plot between Tel1 signals at 3 and 4 h in SPM (log of normalised count over 1500pb regions centered on hotspots), and DSB signals (Spo11-oligos – log of HpM: hits per million mapped reads).

**c**, Scatter plot between Spo11 signals at 3 and 4 h in SPM (log of normalised count over 1500pb regions centered on hotspots), and DSB signals (Spo11-oligos – log of HpM: hits per million mapped reads).

**d**, Boxplots of Spo11-oligos (HpM) partitioned by DSB strength percentile (Spo11-oligos HpM). **e**, Table of *Pearson* correlation on the top 20% hotspots (n=296) between DSBs (Spo11-oligos HpM), Tel1 (normalised count over 1500pb regions centered on hotspots) and Spo11 (normalised count over 1500pb regions centered on hotspots) at 3 and 4 h in SPM, relative to each other’s.

**f**,**g**, Scatter plot between Spo11 and Tel1 signals (log of normalised count over 1500pb regions centered on top 20% hotspots) at 3 h (**f**) and 4 h (**g**) in SPM. *R²* Indicate the coefficient of determination.

Figure S3

**a,** FLAG-Tel1 (blue) and Spo11-FLAG (orange) raw coverage signal in a *sae2Δ* background. Note that a similar signal is observed in the two pre-meiotic (0h) samples and the untagged *sae2Δ* control strain. “High-chippable” regions are peaks found in pre-meiotic and untagged samples (see Methods - Peak calling/High-ChIPable regions)

**b**, Spatial and quantitative agreement of Tel1 signal (upper panels) with DSB signal determined by CC-seq (lower panels). The ChIP-seq profile is shown as an example on Chr3, but Tel1 and CC-seq signals agree well on all chromosomes. The position of the centromere is indicated with the black circle on the horizontal axis. Signal values are plotted after smoothing with an increasing sliding Hann window size as indicated.

**c**, **d**, Correlation between biological replicates (exp2, 6 h). Tel1 (**c**) and Spo11 (**d**) using the average signals in individual 100-bp non-overlapping genomic windows. Coefficient of determination (R²) and associated p-value were determined using linear model fitting function (*lm*).

**e**, FLAG-Tel1 read densities (RPKM per 20-bp bins) ±5 kb from DSBs midpoint in a *sae2Δ* background, from exp1 at 0h (blue light line), 4 h (blue medium line) and 6 h (blue dark line) in SPM. Dashed black line represent the profile of non-specific signal (no-tag).

**f**, FLAG-Tel1 read densities (RPKM per 20-bp bins) ±4 kb from the top 20% hotspots midpoint, in a *sae2Δ* background at 4h (blue light line) versus WT background at 4h (blue dark line).

Figure S4

**a,** Venn diagrams showing the overlap between Hop1, Rec8 and Red1 peak coordinates(53). We define axis coordinates as the intersection of at least two of the three marks (Fig. 2c-e). Percentages of overlap are indicated in the table below.

**b**, Table of overlap between axis, DSBs, Tel1 and Spo11 peaks relative to each other’s (corresponding to percentages in Fig. 2c). Numbers of identified peaks are indicated for the features in the first line relative to features in the first column. “Alone” in the bottom line refers to non-overlapping peaks with any features.

**d**, **e**, Overlap between axis, DSBs and Tel1 peaks. Boxplots are comparing the overlap between real and shuffled distribution of Tel1 (**c**) or DSBs (**d**) peaks along the genome with the indicated feature. Ten independent random genomic region shuffling were generated. Centre, lower edge, and upper edge of the boxplot denote median, 25th, and 75th percentiles; lower end and higher end of the vertical lines denote 5th and 95th percentiles. Mann Whitney test p-values are indicated.

Figure S5

Boxplot of Tel1, Spo11 and DSBs peak strength distribution. Tel1 and Spo11 peak strength was quantified by RPKM (see methods). DSBs peak strength was quantified by NormHpM (methods). The solid horizontal line represents the median, and the box encompasses the lower and upper quartiles. The vertical lines denote 5th and 95th percentiles. Mann Whitney test p-values are indicated.

**a**, Tel1 peaks are distributed according to their overlap with Spo11, DSBs, axis, or with none of the above (‘alone’). Tel1 peaks not associated with other features (Spo11, DSBs or axis sites) are highly significantly weaker than all the others.

**b**, DSBs peaks are distributed according to their overlap with Tel1, Spo11, axis, or with none of the above (‘alone’). DSBs peaks associated with axis sites, or not associated with other features (Tel1, Spo11 or axis sites) are significantly weaker than all others.

Figure S6

**a,** Immunoblot of meiotic TCA extracts at indicated time-points probed with anti-FLAG antibody (Tel1). Bottom panels: anti-RAP1 antibody.

**b**, Quantification of immunoblots (n=3) as per (a). Signal normalization was performed against RAP1 signal.

**c**, FLAG-Tel1-*ΔC* recruitment to meiotic hotspots measured by ChIP-qPCR at the *ARE1, ATF2 and GAT1* hotspot relative to three control sites (CDC39, DRS2 and YCR013C) in *sae2Δ ndt80Δ* and *sae2Δ ndt80Δ FLAG-tel1- ΔC* strains. Error bars indicate SD from 3 (t 0 h) and 5 (t 6 h) independent experiments. P values are from paired ANOVA test between time 0 h and 6 h, corrected via Two-stage linear step-up procedure of Benjamini, Krieger and Yekutieli for multiple comparisons test. ns (not significant), ^∗∗∗∗^, p < 0.0001.

Figure S7

**a,** Boxplot of DSBs, Tel1 and Spo11 normalized counts per 600bp (see Methods) partitioned by DSB strength percentile (NormHPM).

**b**, Boxplot of Axis (Hop1), Tel1 and Spo11 normalized counts per 2kb (see Methods) partitioned by Hop1 strength percentile (RPKM).

Mann Whitney test p-values are indicated.

Figure S8

Heatmap of CC-seq (DSBs) and ChIP-seq read densities of the indicated features ±5 kb from DSBs centres (**a**) or axis centres (**b**). Regions are ordered according to DSBs strength (NormHpM) in all heatmaps presented in (**a**) or according to Hop1 ChIP-seq signal(53) (RPKM) in all heatmaps presented in (**b**).

Figure S9

**a,** Diagram of the *MAT* (HO site) with the primer pairs to measure Tel1 binding by ChIP-qPCR.

**b**, Tel1 recruitment to the Gal::HO break is dependent of the C-terminal part of Xrs2. Recruitment of FLAG-Tel1 was measured by ChIP-qPCR at 0’, 90’ and 180’ after induction of HO expression. Recruitment was measured at 0.15 kb (blue) and 10 kb (red) of the break. Bars represent the mean values of three independent experiments with error bars denoting s.d.

Figure S10

Middle-range DSB interference is abolished in the *xrs2-11* mutant.

**a**, Genomic DNA isolated from the indicated time points after induction of meiosis was prepared in plugs and fractionated by PFGE, transferred to nylon membrane and hybridized with a *FRM2* probe located between the *HIS4::LEU2* and *leu2::hisG* hotspots on the left arm of chromosome III. U, uncut parental DNA. Major double-cut signal is indicated by an asterisk. **b**, Quantification of observed double-cut signals in a

Figure S11.

Loss of DSB interference in the *tel1-ΔC* mutant. The *tel1-ΔC* mutation introduces a termination codon at the amino acid residue 2778, removing the amino acids 2778 to 2787.

**a**, **b**, PstI-digested (**a**) or undigested (**b**) genomic DNA isolated from the indicated time points after induction of meiosis was fractionated by electrophoresis, transferred to nylon membrane and hybridized with probes as indicated (Right probe: *MRX2*, Central probe: *LEU2*). P, PstI-digested parental DNA; U, uncut parental DNA. DSB signals are marked with open triangles. Samples are parts of the same membrane that was cropped for presentation purpose.

**c**, Quantification of DSBs and double-cut signals in a,b

**d**, Interference values (I=1-(observedDC/expectedDC), see Methods)

Table S1

List of yeast strains described in this work.

Table S2

List of PCR primers used in this work.

## Notes

### Competing Interest Statement

The authors have declared no competing interest.

### Summary of Updates

Several experiments were performed in this novel version that significantly improve the manuscript and reinforces our conclusions: Tel1 ChIP-seq during meiotic time-courses in a wild-type background, repeat of Tel1 ChIP-seq in a resection deficient background (sae2D) to include additional time-points. Because the mutants used in our study (spo11, xrs2 and tel1) are checkpoint deficient, we also repeated all our time-courses and Tel1 ChIP-qPCR experiments in the ndt80D background (as was already performed for one of the mutants) to prevent precocious cell cycle exit, as requested by the reviewers. We have also tested Tel1 recruitment to an HO-induced DSB to determine whether xrs2 mutants had the same impact on Tel1 recruitment to hyper-resected DSBs in vegetative cells. Conclusions remain unchanged.

