## Supplementary Figures and Table for "Tel1 is recruited at chromosomal loop/axis contact sites to regulate meiotic DNA double-strand breaks interference"

Figure S1

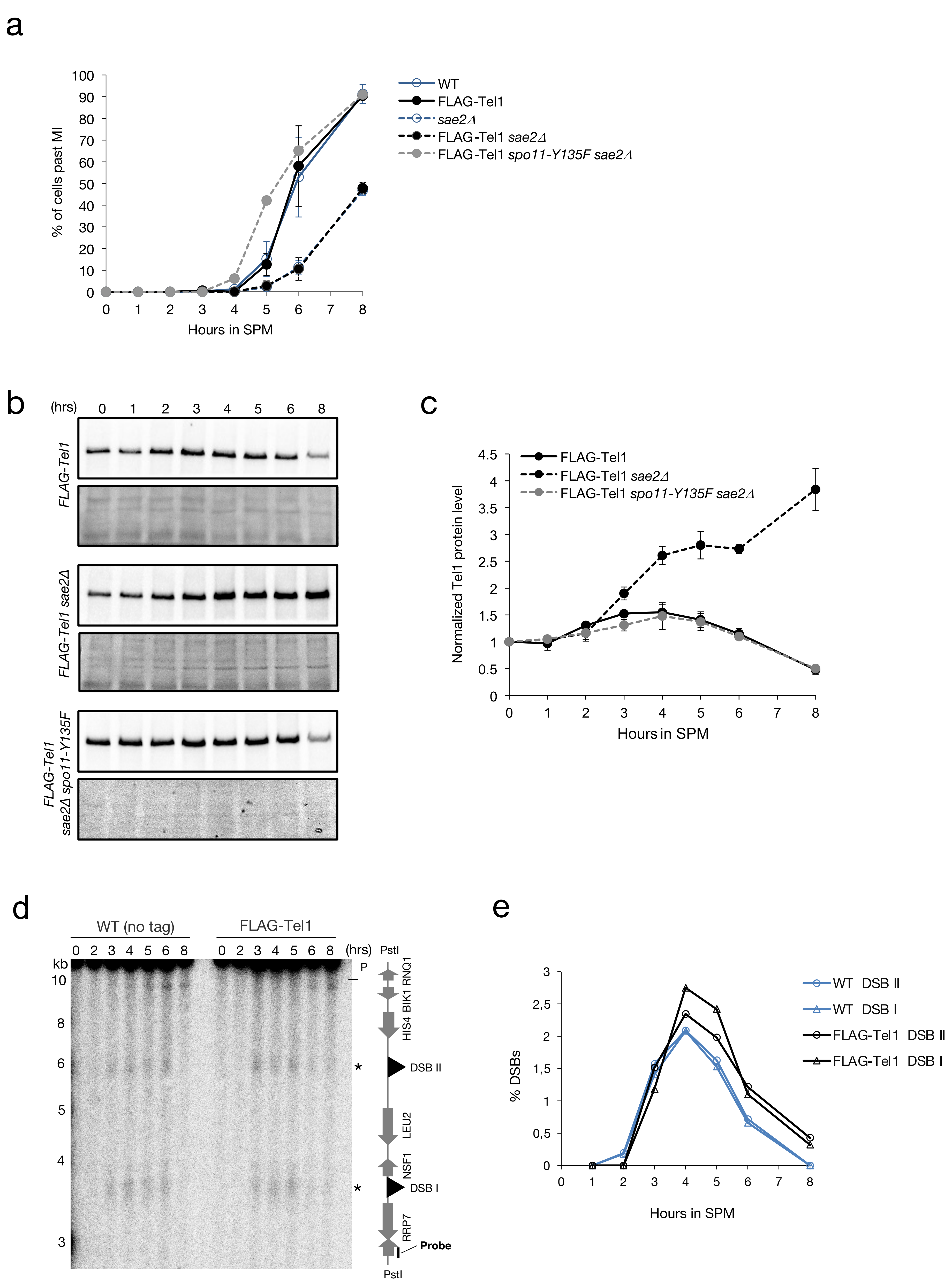

a
% of cells past MI
Hours in SPM
WT
FLAG-Tel1
sae2Δ
FLAG-Tel1 sae2Δ
FLAG-Tel1 spo11-Y135F sae2Δ
b
(hrs)
FLAG-Tel1
FLAG-Tel1 sae2Δ
FLAG-Tel1 sae2Δ spo11-Y135F
c
Normalized Tel1 protein level
Hours in SPM
FLAG-Tel1
FLAG-Tel1 sae2Δ
FLAG-Tel1 spo11-Y135F sae2Δ
d
WT (no tag)
FLAG-Tel1
(hrs)
kb
PstI
P
RNQ1
BIK1
HIS4
DSB II
LEU2
NSF1
DSB I
RRP7
Probe
e
% DSBs
Hours in SPM
WT DSB II
WT DSB I
FLAG-Tel1 DSB II
FLAG-Tel1 DSB I

Figure S2

a

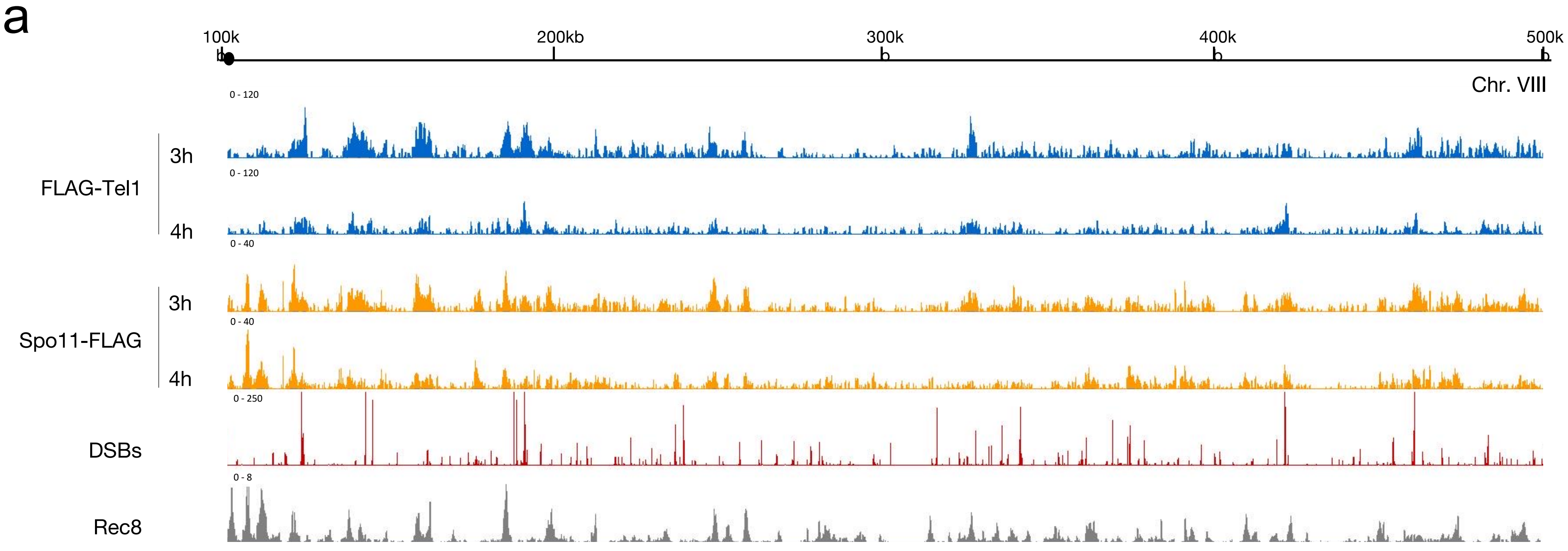


b

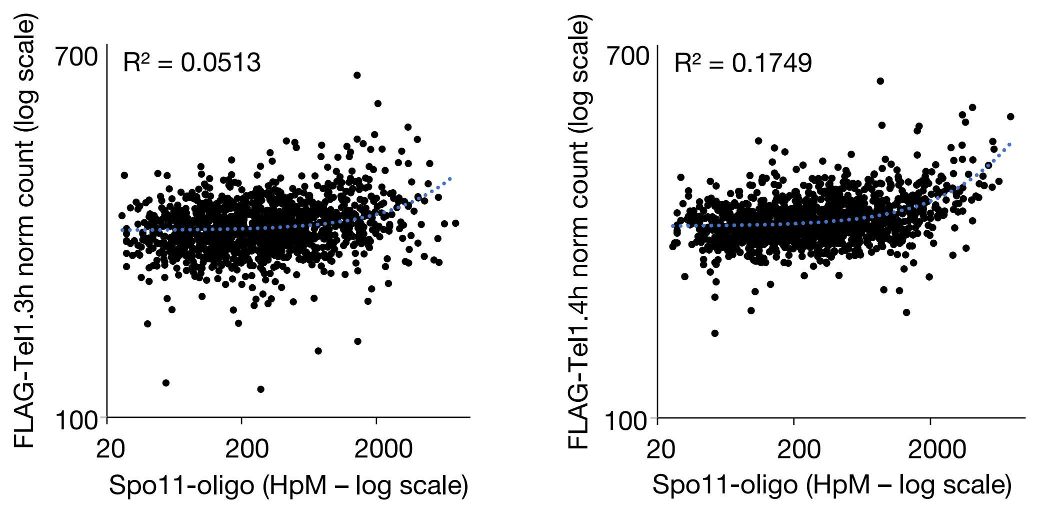


c

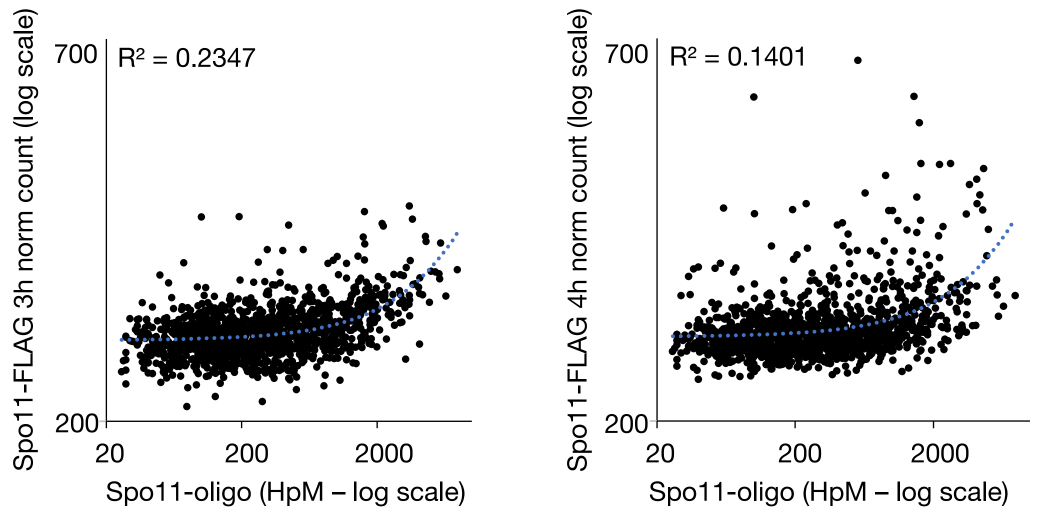


d

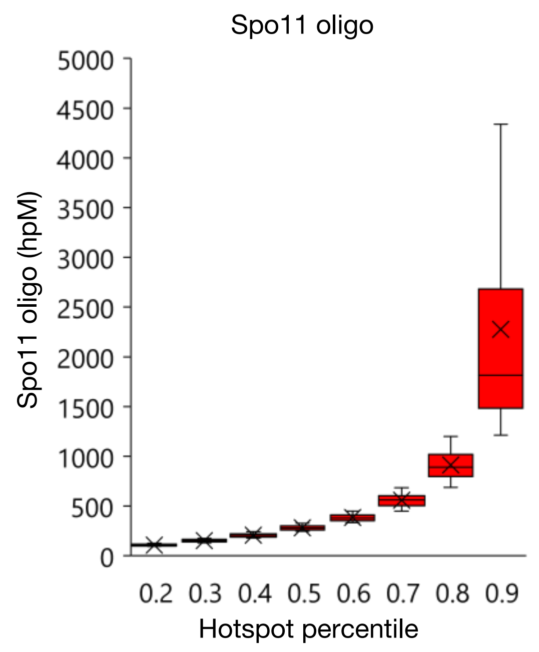


e

| *Pearson top 20% hotspots* | DSB strength | FLAG-Tel1 3h | FLAG-Tel1 4h | Spo11-FLAG 3h | Spo11-FLAG 4h |
|---|---|---|---|---|---|
| DSB strength | 1.00 | 0.42 | 0.43 | 0.48 | 0.30 |
| FLAG-Tel1 3h | 0.42 | 1.00 | 0.56 | 0.37 | 0.09 |
| FLAG-Tel1 4h | 0.43 | 0.56 | 1.00 | 0.46 | 0.24 |
| Spo11-FLAG 3h | 0.48 | 0.37 | 0.46 | 1.00 | 0.61 |
| Spo11-FLAG 4h | 0.30 | 0.09 | 0.24 | 0.61 | 1.00 |

f

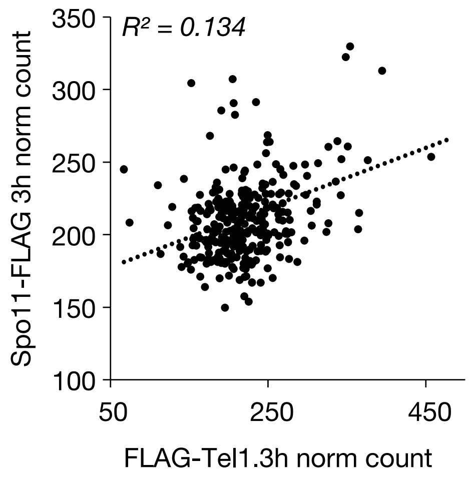


g

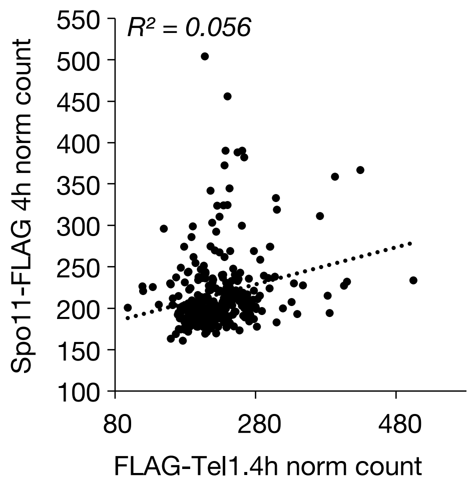

Figure S3

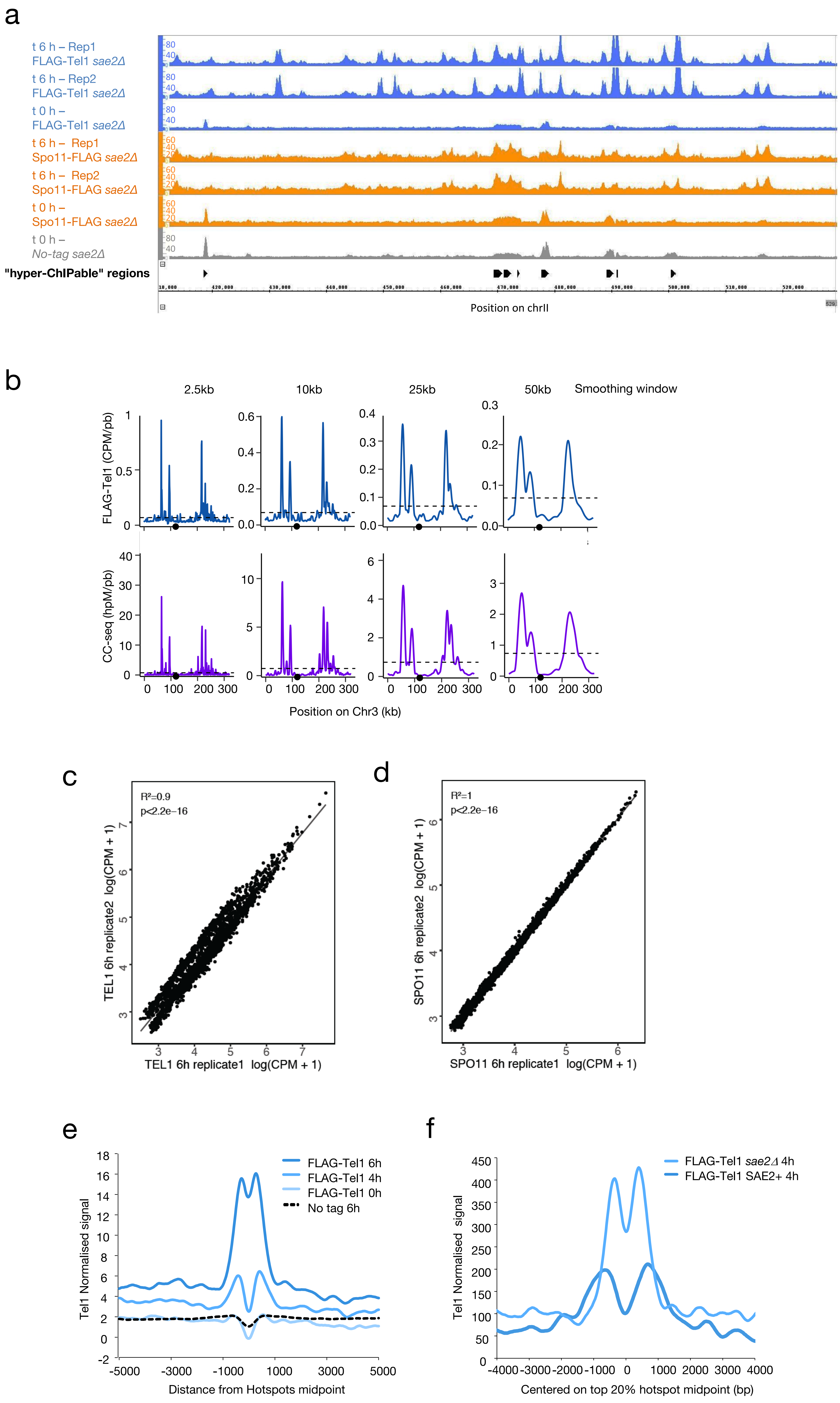

a
t 6 h – Rep1
FLAG-Tel1 sae2Δ
t 6 h – Rep2
FLAG-Tel1 sae2Δ
t 0 h –
FLAG-Tel1 sae2Δ
t 6 h – Rep1
Spo11-FLAG sae2Δ
t 6 h – Rep2
Spo11-FLAG sae2Δ
t 0 h –
Spo11-FLAG sae2Δ
t 0 h –
No-tag sae2Δ
"hyper-ChIPable" regions
Position on chrII
b
2.5kb
10kb
25kb
50kb
Smoothing window
FLAG-Tel1 (CPM/pb)
CC-seq (hpM/pb)
Position on Chr3 (kb)
c
R²=0.9
p<2.2e-16
TEL1 6h replicate2 log(CPM + 1)
TEL1 6h replicate1 log(CPM + 1)
d
R²=1
p<2.2e-16
SPO11 6h replicate2 log(CPM + 1)
SPO11 6h replicate1 log(CPM + 1)
e
FLAG-Tel1 6h
FLAG-Tel1 4h
FLAG-Tel1 0h
No tag 6h
Tel1 Normalised signal
Distance from Hotspots midpoint
f
FLAG-Tel1 sae2Δ 4h
FLAG-Tel1 SAE2+ 4h
Tel1 Normalised signal
Centered on top 20% hotspot midpoint (bp)

Figure S4

a

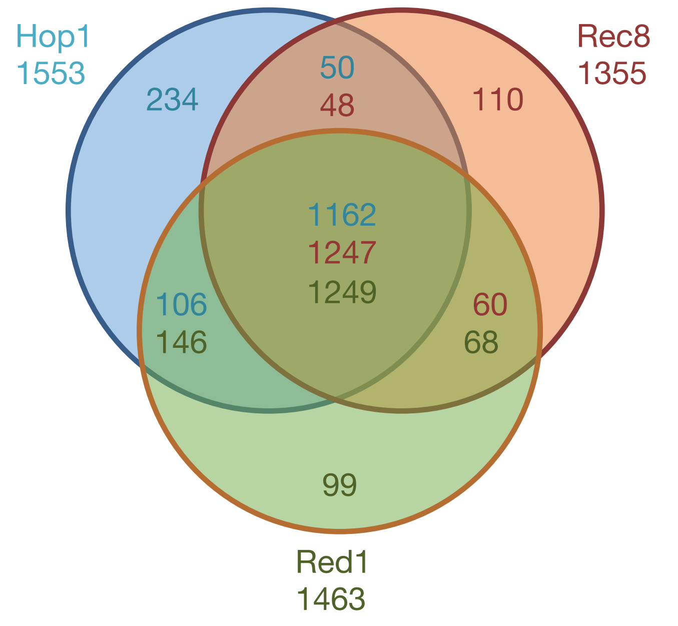


| | UNIQUE | "Hop1" and "Rec8" | "Hop1" and "Red1" | "Rec8" and "Red1" | "Hop1", "Rec8" and "Red1" |
|---|---|---|---|---|---|
| Hop1 | 15.1% | 3.2% | 6.8% | | 74.9% |
| Rec8 | 7.5% | 3.3% | | 4.1% | 85.1% |
| Red1 | 6.3% | | 9.3% | 4.4% | 80.0% |

b

| Sites | Axis | DSB | Tel1 | Spo11 |
|---|---|---|---|---|
| **Axis** | **1357** | 789 | 1468 | 1427 |
| **DSB** | 535 | **3601** | 1818 | 1681 |
| **Tel1** | 1032 | 2143 | **3103** | 2126 |
| **Spo11** | 1177 | 2162 | 2504 | **3042** |
| **alone** | 114 | 910 | 230 | 489 |

c

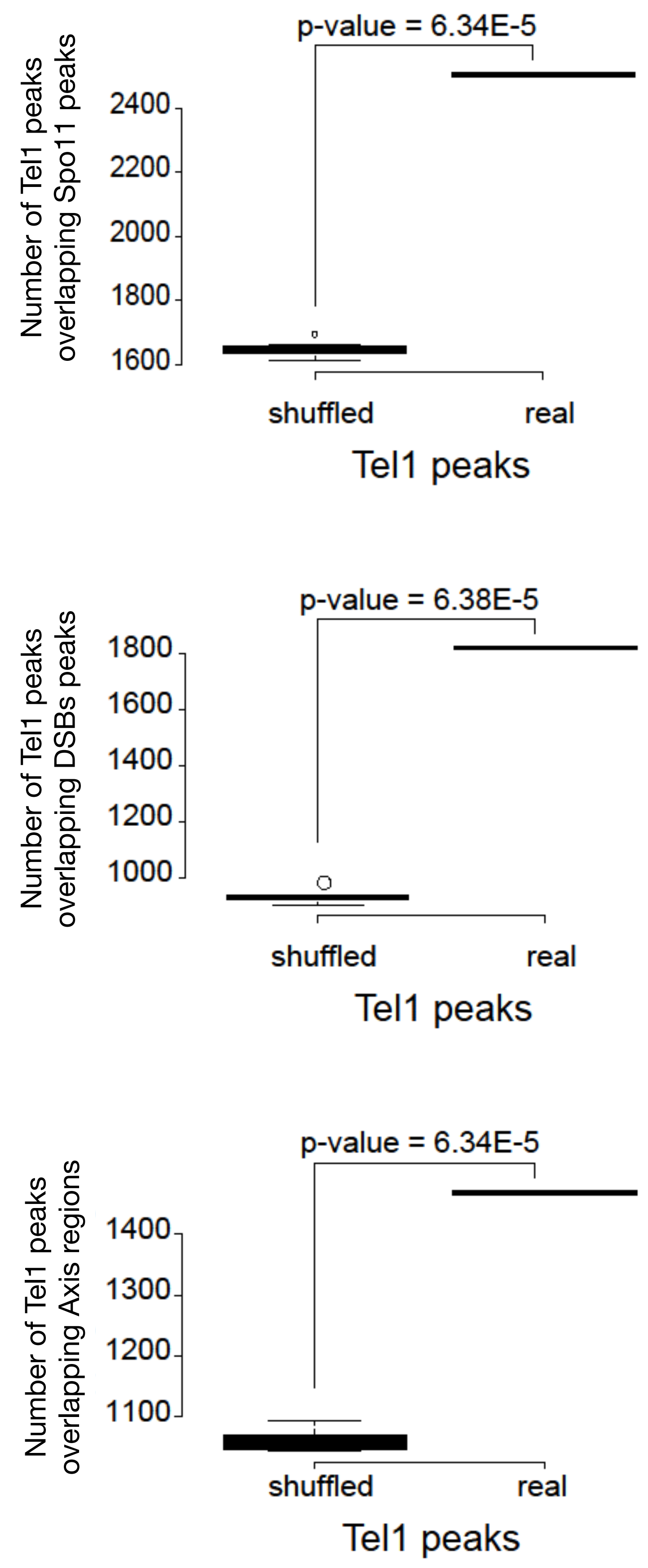


d

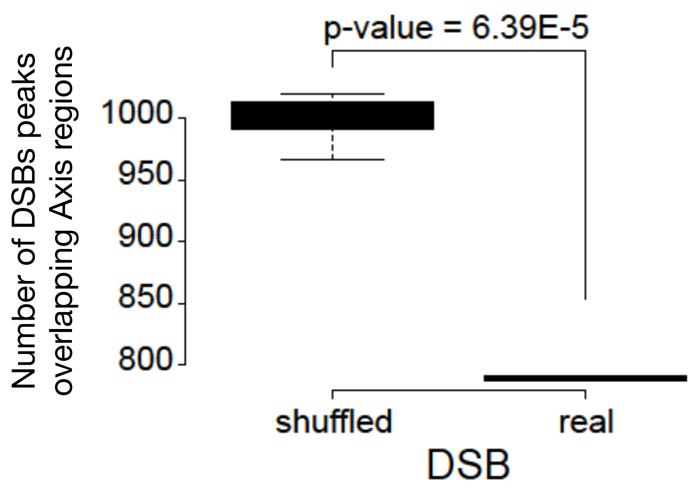

Figure S5

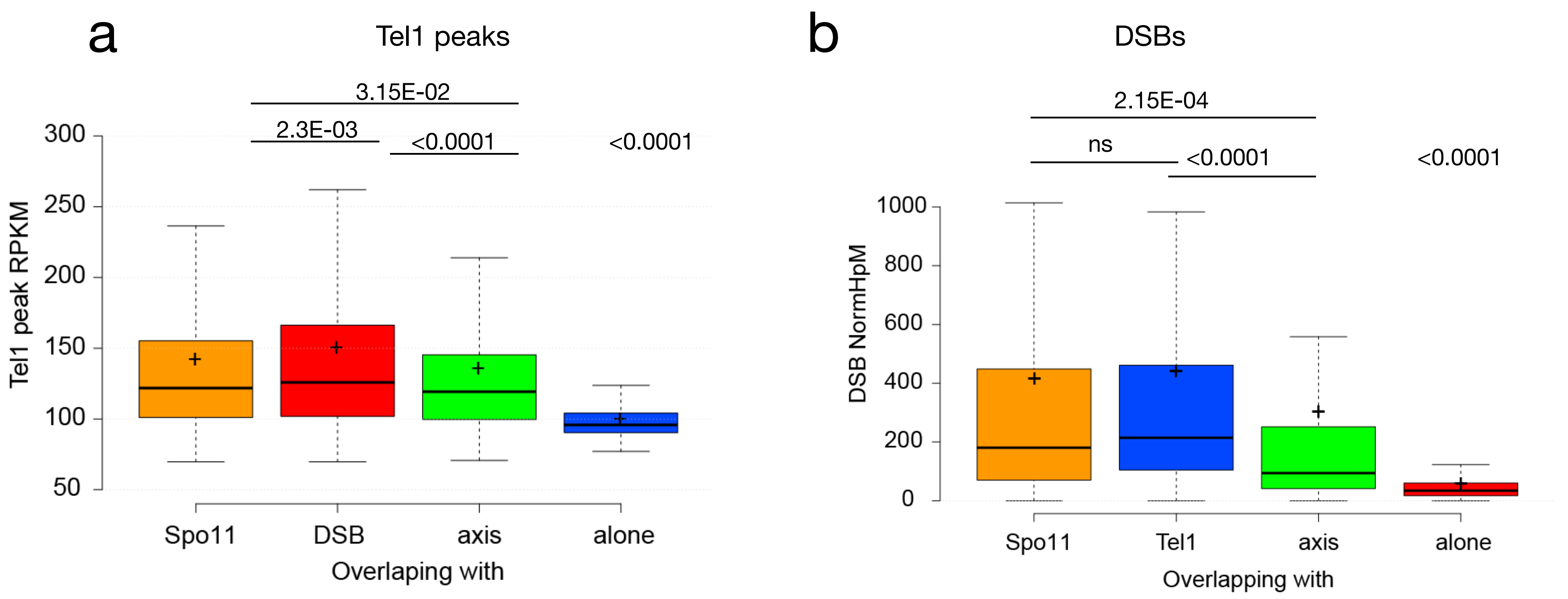

a
Tel1 peaks
3.15E-02
2.3E-03
<0.0001
<0.0001
Tel1 peak RPKM
300
250
200
150
100
50
Spo11
DSB
axis
alone
Overlaping with
b
DSBs
2.15E-04
ns
<0.0001
<0.0001
DSB NormHpM
1000
800
600
400
200
0
Spo11
Tel1
axis
alone
Overlapping with

Figure S6

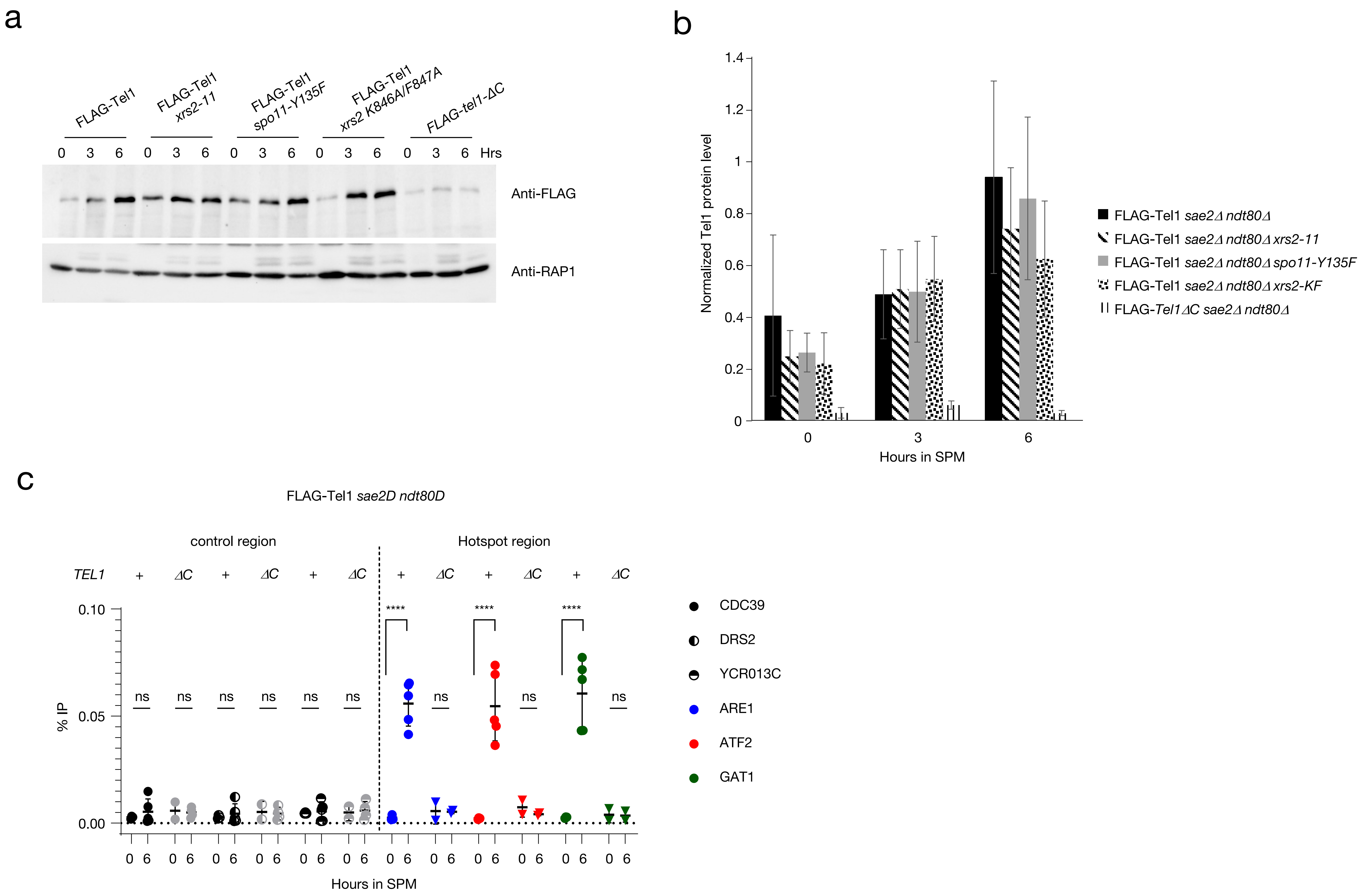
a
FLAG-Tel1
FLAG-Tel1 xrs2-11
FLAG-Tel1 spo11-Y135F
FLAG-Tel1 xrs2 K846A/F847A
FLAG-tel1-ΔC
0 3 6 0 3 6 0 3 6 0 3 6 0 3 6 Hrs
Anti-FLAG
Anti-RAP1
b
Normalized Tel1 protein level
1.4
1.2
1
0.8
0.6
0.4
0.2
0
0
3
6
Hours in SPM
FLAG-Tel1 sae2Δ ndt80Δ
FLAG-Tel1 sae2Δ ndt80Δ xrs2-11
FLAG-Tel1 sae2Δ ndt80Δ spo11-Y135F
FLAG-Tel1 sae2Δ ndt80Δ xrs2-KF
FLAG-Tel1ΔC sae2Δ ndt80Δ
c
FLAG-Tel1 sae2D ndt80D
control region
Hotspot region
TEL1 + ΔC + ΔC + ΔC + ΔC + ΔC + ΔC
0.10
0.05
0.00
% IP
ns
****
0 6 0 6 0 6 0 6 0 6 0 6 0 6 0 6 0 6 0 6 0 6 0 6
Hours in SPM
CDC39
DRS2
YCR013C
ARE1
ATF2
GAT1

Figure S7

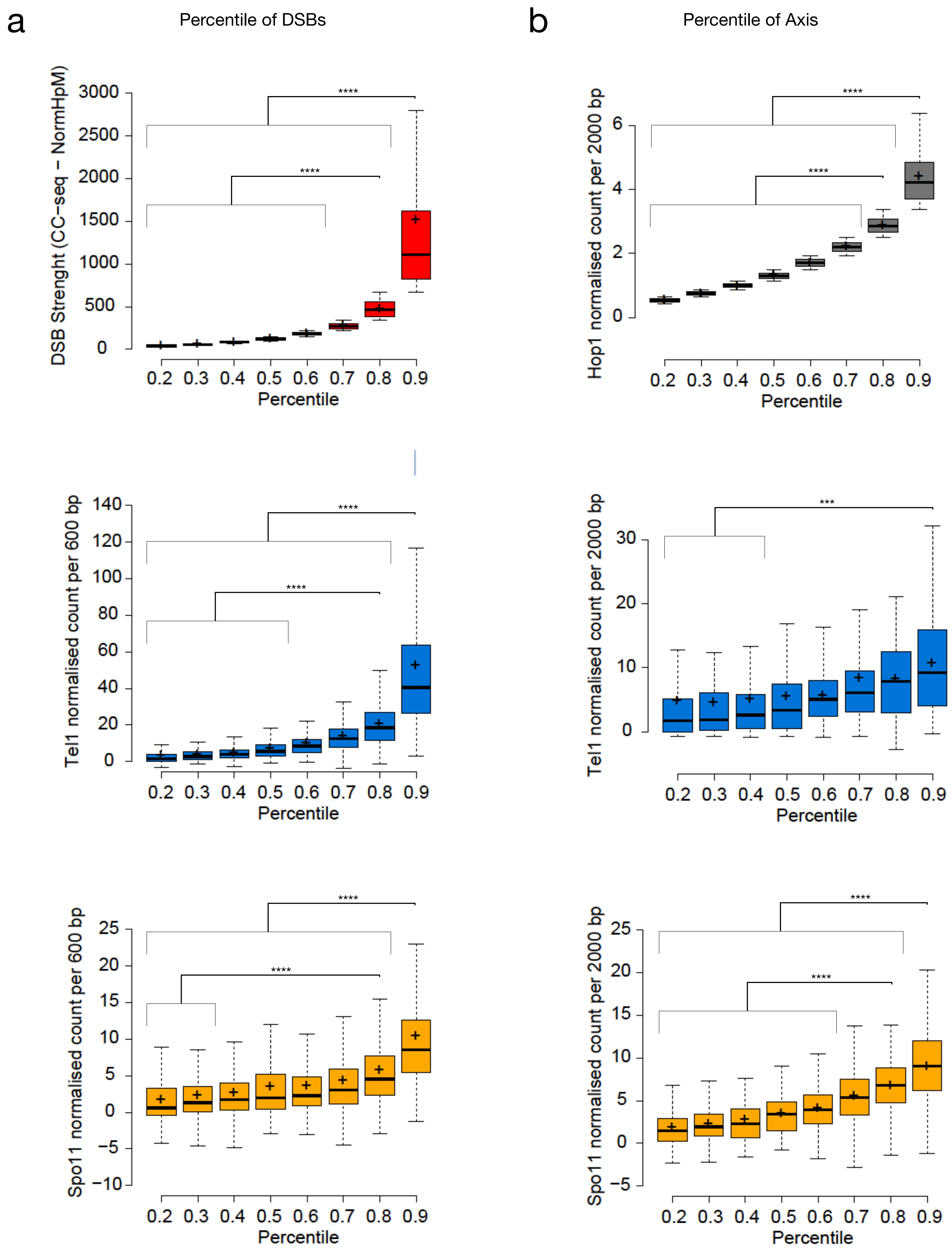
a
Percentile of DSBs
DSB Strenght (CC−seq − NormHpM)
Tel1 normalised count per 600 bp
Spo11 normalised count per 600 bp
Percentile
****
b
Percentile of Axis
Hop1 normalised count per 2000 bp
Tel1 normalised count per 2000 bp
Spo11 normalised count per 2000 bp
Percentile
****
***

Figure S8

a

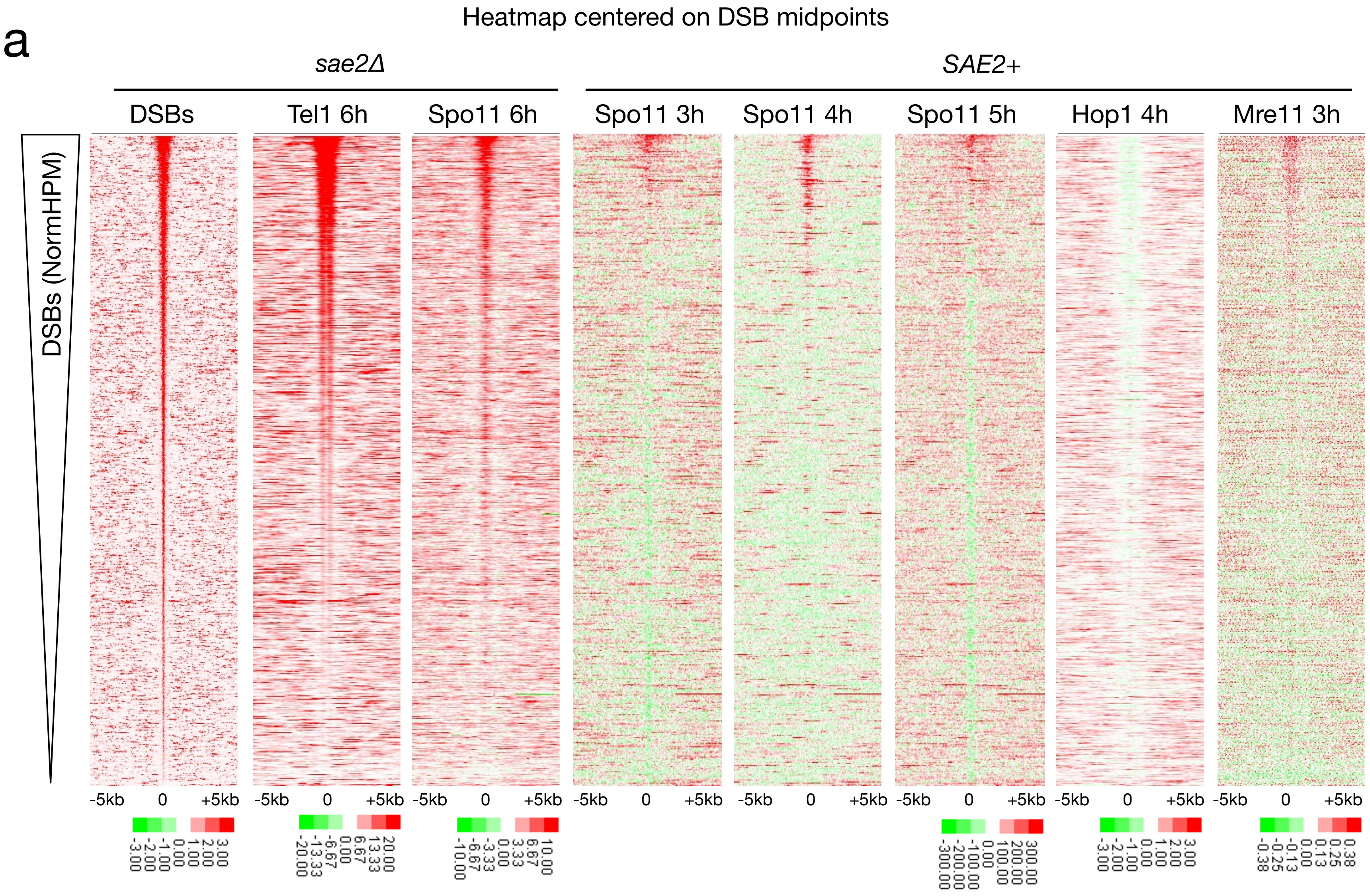

Heatmap centered on DSB midpoints
sae2Δ
SAE2+
DSBs
Tel1 6h
Spo11 6h
Spo11 3h
Spo11 4h
Spo11 5h
Hop1 4h
Mre11 3h
DSBs (NormHPM)
-5kb
0
+5kb


b

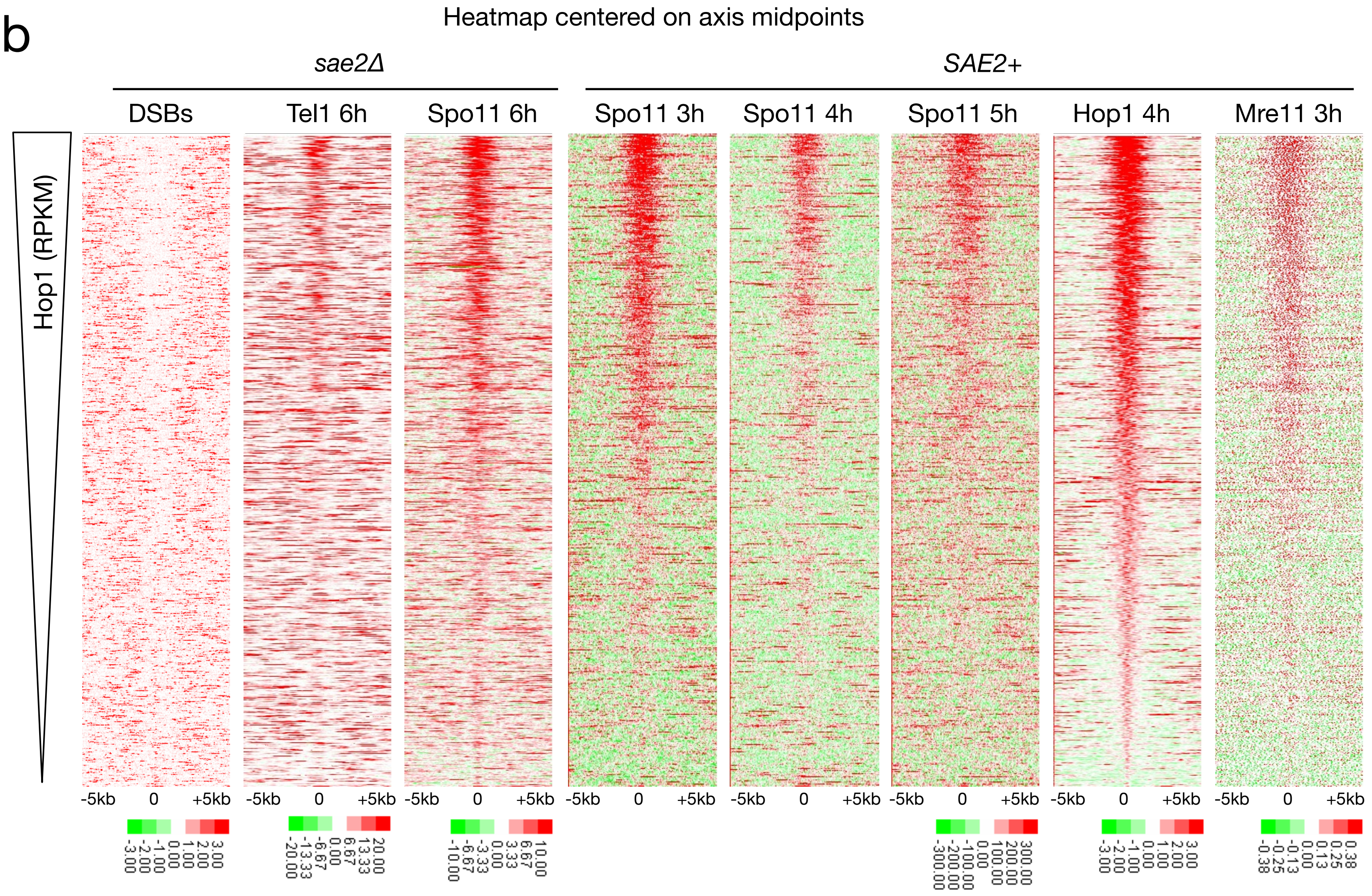

Heatmap centered on axis midpoints
sae2Δ
SAE2+
DSBs
Tel1 6h
Spo11 6h
Spo11 3h
Spo11 4h
Spo11 5h
Hop1 4h
Mre11 3h
Hop1 (RPKM)
-5kb
0
+5kb

Figure S9

a

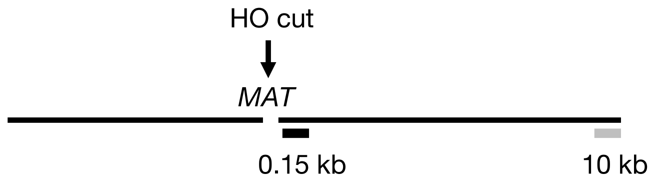

HO cut
MAT
0.15 kb
10 kb


b

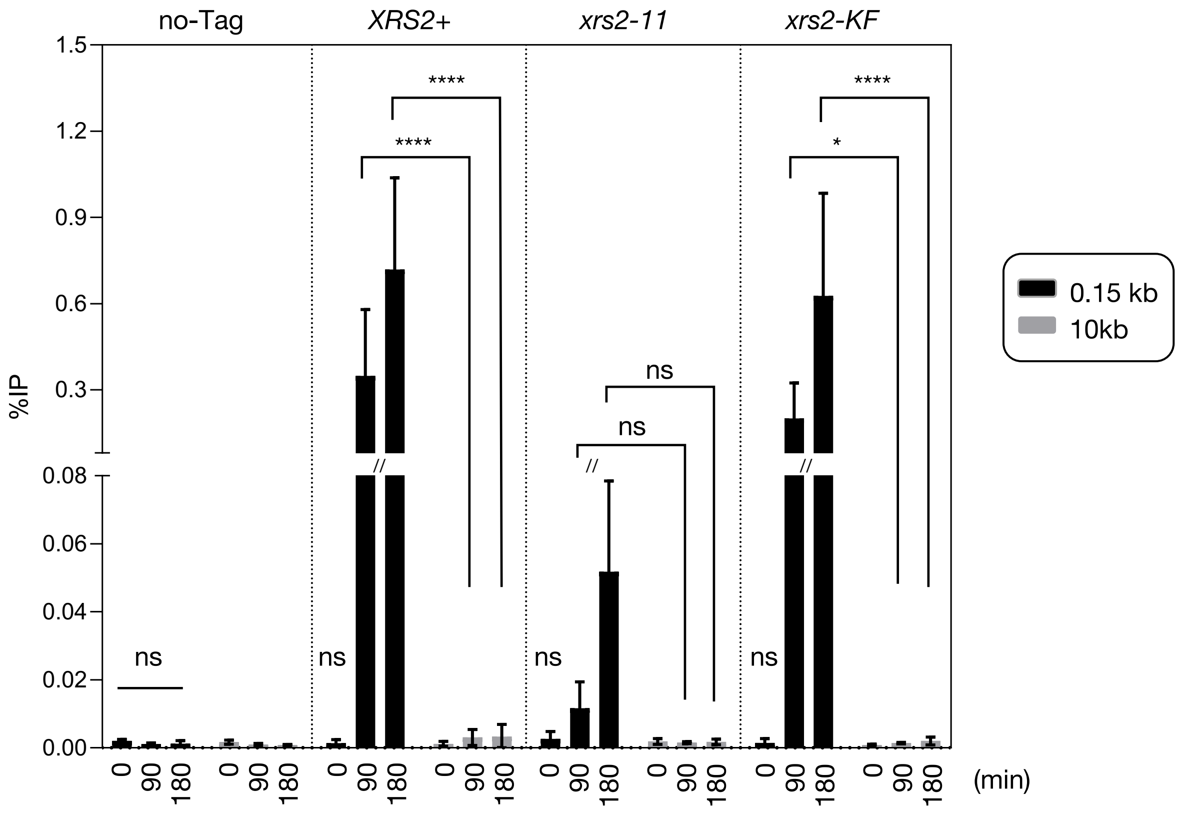

no-Tag
XRS2+
xrs2-11
xrs2-KF
%IP
1.5
1.2
0.9
0.6
0.3
0.08
0.06
0.04
0.02
0.00
ns
****
*
0
90
180
(min)
0.15 kb
10kb

Figure S10

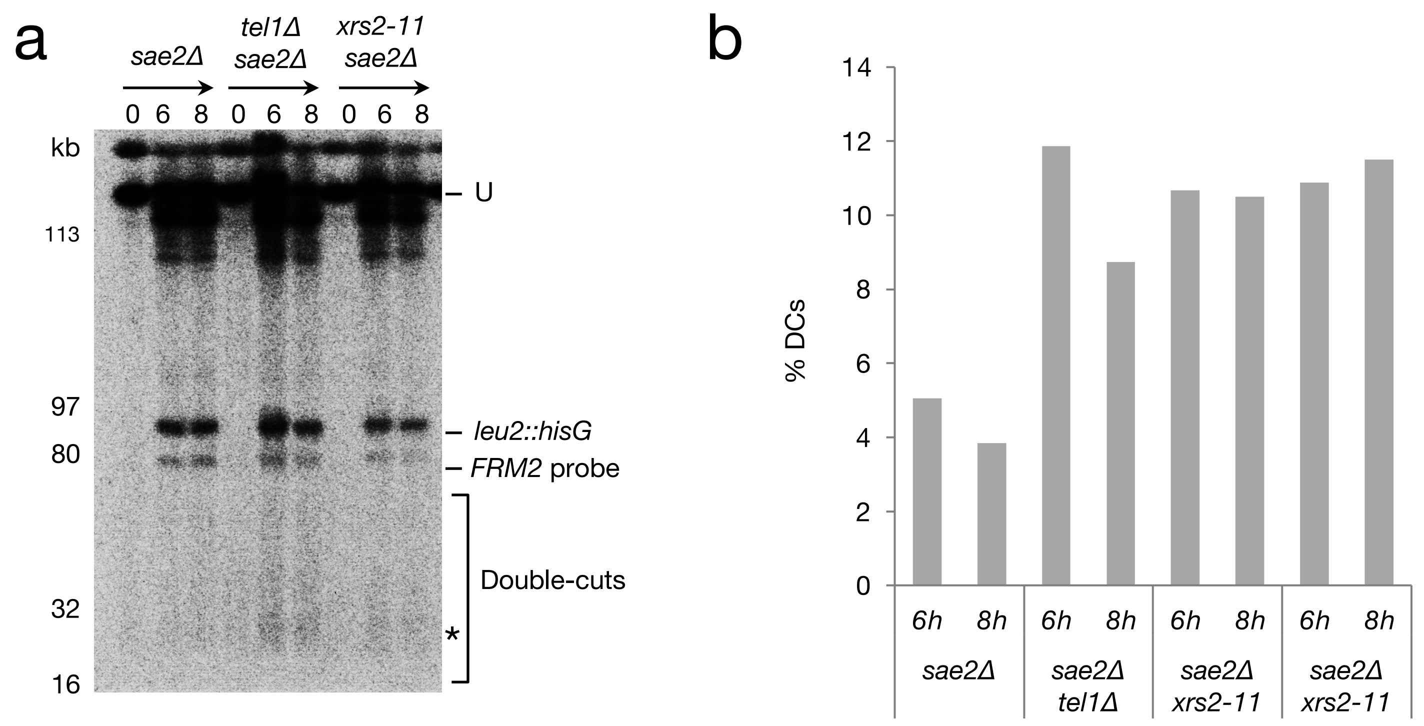
a
sae2Δ
tel1Δ sae2Δ
xrs2-11 sae2Δ
0 6 8 0 6 8 0 6 8
kb
113
97
80
32
16
U
leu2::hisG
FRM2 probe
Double-cuts
*
b
% DCs
14
12
10
8
6
4
2
0
6h 8h 6h 8h 6h 8h 6h 8h
sae2Δ
sae2Δ tel1Δ
sae2Δ xrs2-11
sae2Δ xrs2-11

Figure S11

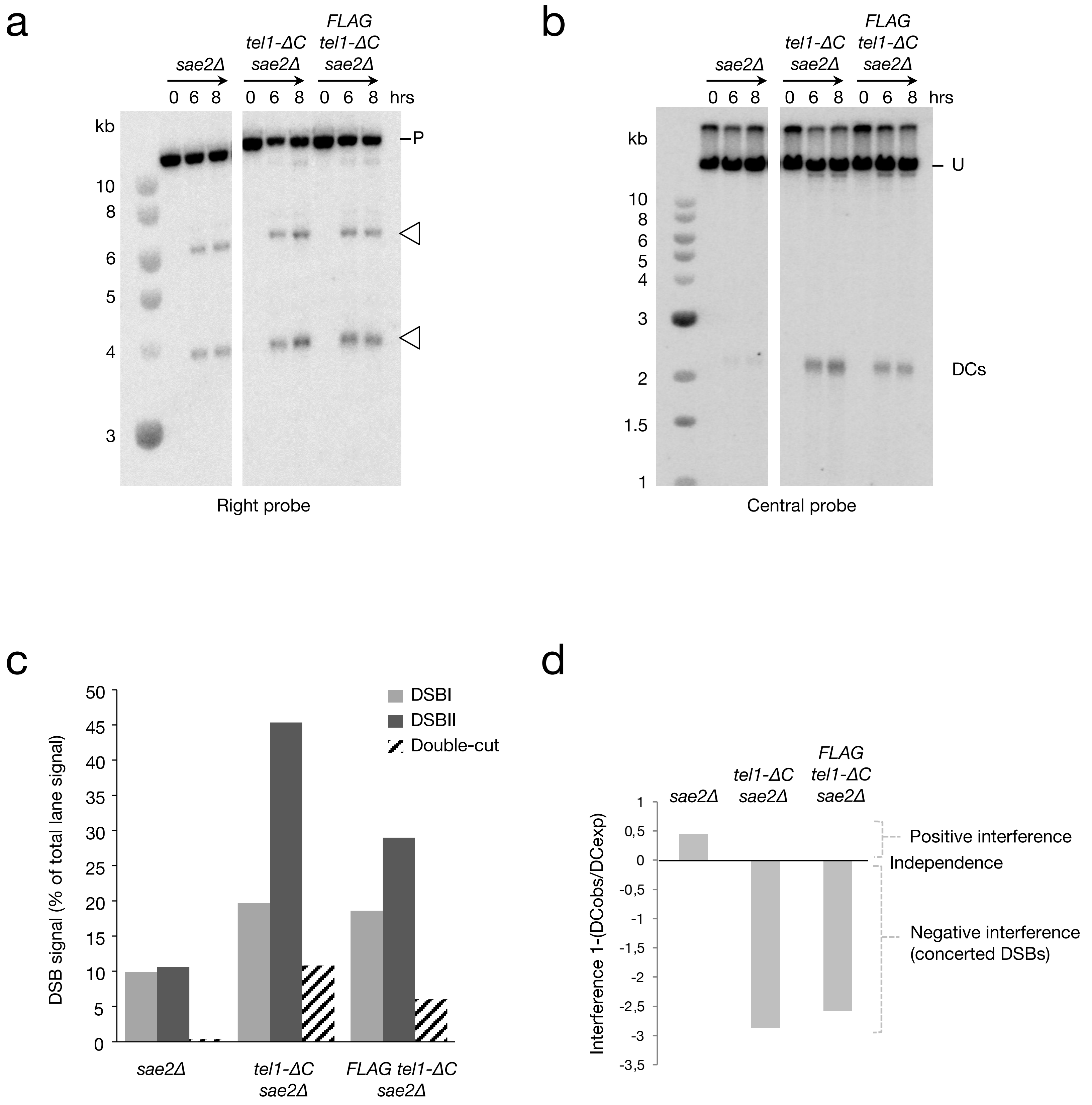
a
sae2Δ
tel1-ΔC sae2Δ
FLAG tel1-ΔC sae2Δ
0 6 8 0 6 8 0 6 8 hrs
kb
10
8
6
5
4
3
P
Right probe
b
sae2Δ
tel1-ΔC sae2Δ
FLAG tel1-ΔC sae2Δ
0 6 8 0 6 8 0 6 8 hrs
kb
10
8
6
5
4
3
2
1.5
1
U
DCs
Central probe
c
DSB signal (% of total lane signal)
50
45
40
35
30
25
20
15
10
5
0
DSBI
DSBII
Double-cut
sae2Δ
tel1-ΔC sae2Δ
FLAG tel1-ΔC sae2Δ
d
Interference 1-(DCobs/DCexp)
1
0,5
0
-0,5
-1
-1,5
-2
-2,5
-3
-3,5
sae2Δ
tel1-ΔC sae2Δ
FLAG tel1-ΔC sae2Δ
Positive interference
Independence
Negative interference (concerted DSBs)

Table S1

| Strain | Mating type | Genotype |
|---|---|---|
| VG499 | a/α | *ho::LYS2/”, lys2/”, ura3/”, arg4-nsp/”, leu2::hisG/”, his4X::LEU2/”, nuc1::LEU2/”, sae2Δ::KanMX6/”* |
| VG739 | a/α | *ho::LYS2/", lys2/", ura3/", arg4-nsp/", leu2::hisG/", his4X::LEU2/", nuc1::LEU2/"* |
| VG740 | a/α | *ho::LYS2/", lys2/", ura3/", arg4-nsp/", leu2::hisG/", his4X::LEU2/", nuc1::LEU2/", FLAG-TEL1", sae2Δ::KanMX6"* |
| VG741 | a/α | *ho::LYS2/", lys2/", ura3/", arg4-nsp/", leu2::hisG/", his4X::LEU2/", nuc1::LEU2/", FLAG-TEL1"* |
| VG906 | a/α | *ho::LYS2/", lys2/", ura3/", arg4-nsp/3, leu2::hisG/", nuc1::LEU2, SPO11-His6-flag3-loxP-HphMX-loxP/", sae2Δ::KanMX6/"* |
| VG613 | a/α | *ho::LYS2/', lys2/', ura3/', arg4-nsp/', leu2::hisG/', his4X::LEU2/', nuc1::LEU2/', xrs2CterΔ/", sae2Δ::KanMX6/"* |
| VG619 | a/α | *ho::LYS2/', lys2/', ura3/', arg4-nsp/', leu2::hisG/', his4X::LEU2/', nuc1::LEU2/', xrs2 K846A/F847A::KANMX6/", sae2Δ::KanMX6/"* |
| VG749 | a/α | *ho::LYS2/", lys2/", ura3/", arg4-nsp/", leu2::hisG/", his4X::LEU2/", nuc1::LEU2/", FLAG-TEL1/", spo11(Y135F)::KanMX4/", sae2Δ::KanMX6/"* |
| VG764 | a/α | *ho::LYS2/", lys2/", ura3/", arg4-nsp/", leu2::hisG/", his4X::LEU2/", nuc1::LEU2/", FLAG-TEL1/", sae2Δ::KanMX6/", xrs2CterΔ::KanMX6/"* |
| VG838 | a/α | *ho::LYS2/", lys2/", ura3/", arg4-nsp/", leu2::hisG/", his4X::LEU2/", nuc1::LEU2/", FLAG-TEL1/", sae2Δ::KanMX6/", xrs2 K846A/F847A::KANMX6/"* |
| VG948 | a/α | *ho::LYS2/", lys2/", ura3/", arg4-nsp/", leu2::hisG/", his4X::LEU2/", nuc1::LEU2/", FLAG-tel1-ΔC::NatMX6/", sae2Δ::KanMX6/"* |
| VG1212 | a/α | *ho::LYS2/', lys2/', ura3/', arg4-nsp/', leu2::hisG/', his4X::LEU2/', nuc1::LEU2/', FLAG-TEL1/', sae2Δ::KanMX6/', xrs2CterΔ::KanMX6/', ndt80::HphMX/'* |
| VG1238 | a/α | *ho::LYS2/", lys2/", ura3/", arg4-nsp/", leu2::hisG/", his4X::LEU2/", nuc1::LEU2/", FLAG-TEL1/", sae2Δ::KanMX6/", ndt80::HphMX/"* |
| VG1582 | a/α | *ho::LYS2/", lys2/", ura3/", arg4-nsp/", leu2::hisG/", his4X::LEU2/", nuc1::LEU2/", FLAG-TEL1/", sae2Δ::KanMX6/",spo11(Y135F) ::KanMX6/" ndt80::HphMX/"* |
| VG1590 | a/α | *ho::LYS2/", lys2/", ura3/", arg4-nsp/", leu2::hisG/", his4X::LEU2/", nuc1::LEU2/", FLAG-TEL1/", sae2Δ::KanMX6/",xrs2 K846A/F847A ::KanMX6/" ndt80::HphMX/"* |
| VG1597 | a/α | *ho::LYS2/", lys2/", ura3/", arg4-nsp/", leu2::hisG/", his4X::LEU2/", nuc1::LEU2/", FLAG-TEL1/", sae2Δ::KanMX6/",FLAG-tel1-ΔC ::NatMX6/" ndt80::HphMX/"* |
| VG1393 | a | *hoΔ hmlΔ ::ADE1 hmrΔ ::ADE1 ade1-100 leu2,3-112 lys5 trp1::hisG ura3-52 ade3::GAL::HO* |
| VG1447 | a | *hoΔ hmlΔ ::ADE1 hmrΔ ::ADE1 ade1-100 leu2,3-112 lys5 trp1::hisG ura3-52 ade3::GAL::HO FLAG-TEL1 xrs2-11* |
| VG1448 | a | *hoΔ hmlΔ ::ADE1 hmrΔ ::ADE1 ade1-100 leu2,3-112 lys5 trp1::hisG ura3-52 ade3::GAL::HO FLAG-TEL1 xrs2-K846A,F847A* |

Table S2

**Xrs2-11_pFA6_F@2030**
ATCACTAGAAATTATGTTCCGTTAAAAAAATACTCCAAAAAGGATACAACTACAAAATGGGTGA GGCGCGCCACTTCTAAA
**Xrs2-11_pFA6_R@2627**
TATCGAATGATAATGCAAAATATAATTTAATGAAATTGGAAATACTCGGAAAATTTATCAATCGATGAATTCGAG CTC
**Xrs2-K846A/F847A_pFA6_F@2030**
TGATGGCGACGACGACGATGACGACGGTCCG GCGGCTACGTTCAAAAGAAGAAAAGGATA TGA GGCGCGCCACTTCTAAA
**Tel1-ΔC_NatMX6_F**
GTAATGGACTAAGTGTAGAGTCTAGCGTACAAGATTTGATTCAGCAAGCCACGGATCCATCAAATTTGAGTtag tgaattcgcgccacttctaaataagc
**Tel1d-ΔC_NatMX6_R**
CATCGCATTGACTTTGTACATTACTTTTCGTATTTCTATAAACAAAAAAAAGAAGTATAAAGCATCTGCATAGCA Aactggatggcggcgttagtatcga

**Primers qPCR**

| **Locus** | | **Site** | **Forward Primer** | **Reverse Primer** | **Fragment size** |
|---|---|---|---|---|---|
| *CDC39* | ChrIII | Control | CDC39_F@+3863<br>CGCTCCACCGATGACTCAAA | CDC39_R@+4014<br>CGACTGGGATGGCTGATTCA | 151pb |
| *MRE11* | ChrXIII | Control | MRE11_F@+505<br>CCACTAAGTTAGCATTGTACGG | MRE11_R@+610<br>CTTCTCGCATAGTCGGTACTTC | 105pb |
| DRS2 | ChrI | Control | DRS2_F@+1454<br>GTACCCTCCAGGTAAAGGTACG | DRS2_R@+1278<br>ATGCGTAATGCTACTGCAACC | 168bp |
| *ATF2* | ChrVII | Hotspot | ATF2@F+1548<br>GAATGGGAATCGTTCTGCAAGC | ATF2@R+1671<br>CACTGCTTGCCTTTTGTACGAG | 123pb |
| *ARE1* | ChrIII | Hotspot | ARE1_F@-58<br>TCACGCAGGTGGTTGTTCAG | ARE1_R@+63<br>GGAATTGAGGCTGCGGATCTTA | 121pb |
| *GAT1* | ChrVI | Hotspot | GAT1@F+25<br>CGCCCTTCCCCTGTTCTG | GAT1@R+165<br>AAATTCAAGTCCGGGTCGAGG | 138pb |
| *YBP1* | ChrII | Axis | YBP1_F@+1811<br>GGCAGATGCCAAGAAGAGTG | YBP1_R@+1962<br>CGGAAGTTTATCAGGTTCGACTG | 151pb |

| | | | | | |
|---|---|---|---|---|---|
| *GRR1* | ChrX | Axis | GRR1_F@+3435<br>GCGTTCCTGATGCTTCATCC | GRR1_R@+3302<br>CCTGACCAGATGAGGAATCTCC | 133pb |

**Primers probes Southern-blot**

| Probe | | Forward Primer | Reverse Primer |
|---|---|---|---|
| *MRX2* | Chr III | HIS4_F@+5170<br>CGTGAAGTGGAACGATGCCC | HIS4_R@+5493<br>GCAACTGTTTCCAGCCTTCACC |
| *LEU2* | Chr III | LEU2_F<br>ATATACCATTCTAATGTCTGC | LEU2_R<br>AAGGATTTTCTTAACTTCTTCGGCG |
| *FRM2* | Chr III | FRM2_F@+27<br>GCTATTACAAACCGTCGTACCATC | FRM2_R@+645<br>CATCGCTGAGGTATCATTACTTCAT |
